# Structural Biophysics-Guided Computational Design of Semaglutide Analogues to Enhance GLP-1R Activation

**DOI:** 10.1101/2024.11.29.625980

**Authors:** Wei Li

## Abstract

CagriSema is a fixed-dose combination of cagrilintide (an amylin analogue) and semaglutide (a GLP-1 receptor agonist), and is currently an experimental obesity drug developed by Novo Nordisk. In March, 2025, CagriSema underperformed expectations in a Phase III trial, achieving 15.7% weight loss instead of the anticipated 25%, raising concerns about its efficacy and clinical value. Given its chemical composition, the weight-loss efficacy of CagriSema is inextricably linked to the activations of GLP-1R and amylin receptors (AMYRs). With GLP-1R as an example target here, this study employs a structural biophysics-guided computational approach for the design of semaglutide analogues to enhance the activation of its receptor GLP-1R. To fully harness the therapeutic potential of GLP-1R activation, an experimental structural basis (PDB entry 4ZGM) of the GLP-1-GLP-1R interaction is essential for the design of semaglutide analogues, where site-specific missense mutations are engineered into its peptide backbone to establish additional stabilizing interactions with the extracellular domain (ECD) of GLP-1R. Specifically, this study puts forward an automated systemic natural amino acid scanning of the peptide backbone of semaglutide, where PDB entry 4ZGM was used as the structural template for high-throughput structural modeling by Modeller and ligand-receptor binding affinity (K_d_) calculations by Prodigy. To sum up, this article reports a total of 564 computationally designed semaglutide analogues with improved GLP-1R ECD binding affinity. Moreover, this study proposes a concept of interfacial electrostatic scaffold comprising four salt bridges at the binding interface of GLP-1R ECD and semaglutide analogues. Drawing parallels with the continued optimization in the past century history of insulin, this article argues that the interfacial electrostatic scaffold here constitutes a robust framework for continued development of next-generation GLP-1R agonists, enabling more effective therapies for patients with diabetes and/or obesity.

## Introduction

Over one century ago, the discovery of insulin marked a transformative milestone in diabetes treatment, leading from life-saving animal-derived therapies to recombinantly synthetic human insulin and its analogues. These continued advancements revolutionized glycemic control for patients with diabetes of both types [1–5]. From its day one, Novo Nordisk has played a pivotal role in this evolution, including also its recent developments of glucagon-like peptide-1 receptor (GLP-1R) agonist such as semaglutide [6–9]. Since its FDA approval in December of 2017, semaglutide has facilitated once-weekly dosing for type 2 diabetes, significantly enhancing patient compliance and clinical outcomes [10–13]. Beyond type 2 diabetes, semaglutide also demonstrated efficacy in weight regulation and cardiovascular risk reduction [14]. Given this, Novo Nordisk continues to explore novel therapeutic avenues, as evidenced by the development of IcoSema [15–19] and CagriSema [20]. Take CagriSema for example, which is a combination therapy comprising cagrilintide and semaglutide, designed for the treatment of type 2 diabetes and obesity. Recently, a Phase III trial results revealed that CagriSema underperformed relative to expectations, achieving 15.7% weight loss instead of the anticipated 25%, raising concerns regarding its overall efficacy and clinical value.

To improve therapeutic efficacy, it is essential to consider the molecular basis of receptor activation and the structural basis of drug-target interaction. Take GLP-1R for example, which is a class B G-protein-coupled receptor (GPCR), and plays a pivotal role in glucose homeostasis by mediating the pharmacological effects of GLP-1 agonists [21, 22]. Previous studies have already highlighted the extracellular domain (ECD) of GLP-1R as a key factor to initiate ligand binding, receptor activation and subsequent intracellular signal transduction [23–26]. As such, enhancing ligand-ECD interactions represents a reasonable structural strategy to augment the therapeutic potential of GLP-1R agonists by enhancing receptor activation. Therefore, this study aims to design semaglutide analogues with site-specific missense mutations to enhance GLP-1R activation through improved ECD binding affinity [21, 27].

## Materials and Methods

As of March 27, 2025, there is one experimental structure determined by X-ray diffraction of the semaglutide backbone in complex with GLP-1R ECD (<u>PDB ID: 4ZGM</u> [24]), which is used here as the template for high-throughput [28] structural modeling by Modeller [29] and ligand-receptor binding affinity (K_d_) calculations by Prodigy [30, 31].

Specifically, this study conducted an automated systemic amino acid scanning of the peptide backbone of semaglutide (Figure 1). Here, **Modigy** is defined as an abbreviation of Modeller [29] and Prodigy [30, 31] to represent an in silico high-throughput generation of structural and intermolecular binding affinity (K_d_) data [28]. Here, **Modigy** follows three criteria:

**Figure 1.**
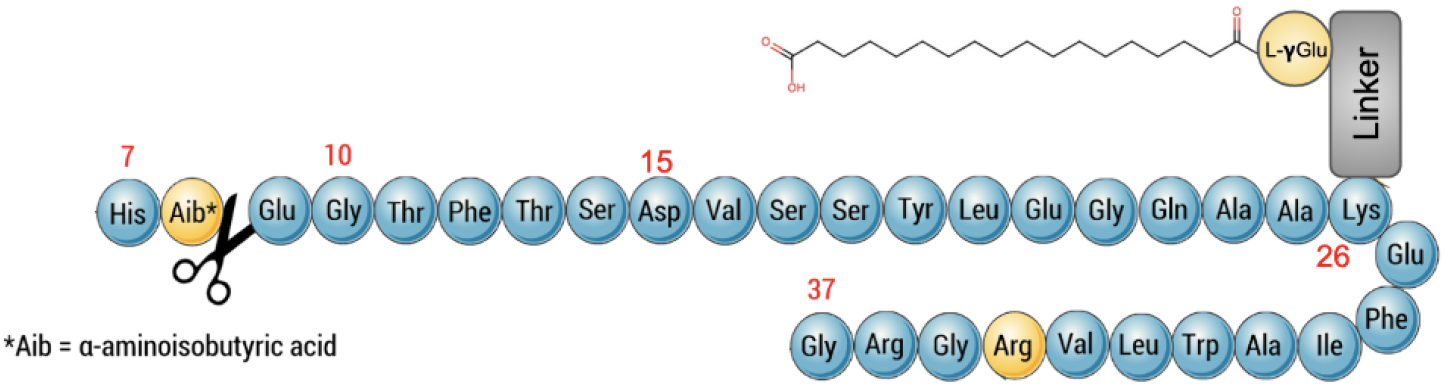
2D structure of semaglutide. In this figure, the sequence of semaglutide’s backbone is numbered from 7 to 37, with modifications highlighted in yellow and the dipeptidyl peptidase-4 (DPP4) cleave site [32, 33] indicated with a scissor. This figure is obtainted from the PDB website (URL: https://pdb101.rcsb.org/global-health/diabetes-mellitus/drugs/incretins/drug/semaglutide/semaglutide, accessed on November 22, 2024).

1. position 8 (Aib near the black scissor in Figure 1) remains unchanged, to avoid DPP4 cleavage [32, 33] of the computationally designed analogues;
2. position 26 (Lys26) of semaglutide’s backbone (Figure 1) remains unchanged, to allow attachment of the fatty acid chain to the peptide backbone;
3. there is just one lysine (i.e., Lys26) in the backbone (Figure 1) of semaglutide, to ensure correct attachment of the fatty acid chain to the peptide backbone of semaglutide.

Subsequently, for all Modeller-generated [29] structural models, a comprehensive structural biophysical analysis [34] was conducted to identify of key electrostatic interactions at the binding interface of GLP-1R and semaglutide analogues [24, 27]. Specifically, the cut-off distance is 4.0 A° for salt bridge analysis [34], while the hydrogen bond analysis employed two criteria: (a) a cutoff value of the angle formed by acceptor (A), donor (D) and hydrogen (H) (∠*ADH*) of 30°; a cutoff value of donor-acceptor (D-A) distance at 3.0 °A [34]. For **Modigy** [28], homology structural modeling by Modeller [29] and K_d_calculations by Prodigy [30, 31] were performed on the high performance computing (HPC) platform at Wuxi Taihu Lake HPC center.

## Results

As mentioned above, semaglutide (GLP-1R agonist) binding directly to GLP-1R ECD [24] is the first step that initiates downstream GLP-1R activation and transmembrane sigal transduction [25, 26]. As such, the primary focus of this study is the interaction between GLP-1R ECD and semaglutide, where PDB entry 4ZGM [35] constitutes the structural basis for subsequent computational design of semaglutide analogues with improved GLP-1R ECD binding affinity.

### A structural biophysical investigation into PDB entry 4ZGM

In 2015, a team of Novo Nordisk deposited into Protein Data Bank [36] the first experimental structure (PDB entry 4ZGM [24, 35]) of semaglutide backbone in complex with the GLP-1R ECD, which was used as the template for **Modigy** [28–31]. Thus, PDB entry 4ZGM was first subject to a comprehensive structural biophysical analysis [34], leading to the identification of two interfacial salt bridges (Table 1) and two interfacial hydrogen bonds (Table 2) at the binding interface of semaglutide and GLP-1R ECD.

**Table 1:** Interfacial salt bridge analysis of PDB entry 4ZGM [24, 35]). In this table, the residue naming scheme is Chain ID Residue Name_Residue ID, and the residue ID numbering scheme is described in Table 1 of supplementary file **supps.pdf**, and chains A and B represent GLP-1R ECD and semaglutide, respectively.

| PDB ID | Residue A | Atom A | Residue B | Atom B | Distance (Å) |
| --- | --- | --- | --- | --- | --- |
| 4ZGM | B_LYS_26 | NZ | A_GLU_128 | OE1 | 3.409 |
| 4ZGM | B_LYS_26 | NZ | A_GLU_128 | OE2 | 2.771 |

**Table 2:**
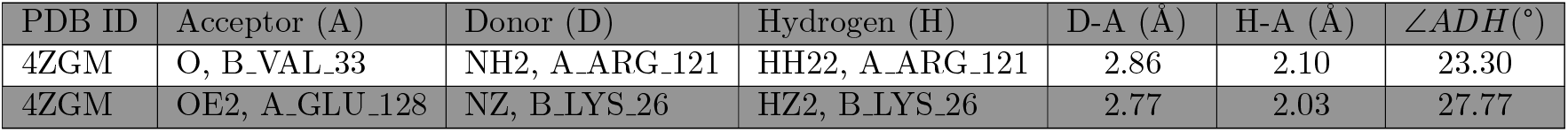
Interfacial side chain hydrogen bond analysis of PDB entry 4ZGM [24, 35]. In this table, the residue naming scheme is the same as Table 1, the residue ID numbering scheme is described in Table 1 of supplementary file **supps.pdf**, while ∠*ADH* represents the angle formed by acceptor (A), donor (D) and hydrogen (H), and chains A and B represent GLP-1R ECD and semaglutide, respectively.

Subsequently, the binding affinity of native semaglutide and GLP-1R ECD was calculated by Prodigy [30, 31] to be 3.4 *×* 10^−6^M at 37 ^°^C, with the template being <u>PDB entry: 4ZGM</u> [24], where a total of 42 residue pairs were found to be located at the binding interface of semaglutide and GLP-1R ECD [24, 35], as listed in Table 4 of supplementary file **supps.pdf**. According to the threshold distance of 5.0 °A used by Prodigy [30, 31], among the 42, only two pairs of charged residues are in interfacial contact: (a) B_LYS_26 of semaglutide and A_GLU_128 of GLP-1R ECD; (b) B_ARG_36 of semaglutide and A_GLU_68 of GLP-1R ECD [24].

As such, the interfacial residue pair of B LYS 26 and A GLU 128 plays an pivotal role both in the binding of semaglutide to GLP-1 ECD and in the stabilization of the ligand-receptor complex structure [21, 37–41], because

1. the two oppositely charged residues are in interfacial contact, according to the Prodigy [30, 31] analysis of PDB entry 4ZGM [24];
2. two interfacial salt bridges (Table 1) formed between the oppositely charged side chains of B_LYS_26 and A_GLU_128;
3. one interfacial hydrogen bond (Table 2) formed between the oppositely charged side chains of B_LYS_26 and A_GLU_128.

Apart from the residue pair of B_LYS_26 and A_GLU_128 (Tables 1 and 2), the Prodigy [30,31] analysis of PDB entry 4ZGM [24] identified another interfacial residue pair of B ARG 36 of semaglutide and A_GLU_68 of GLP-1R ECD [24], for which a detailed distance analysis is included in Table 3. For B_ARG_36 and A_GLU_68, the distances between their oppositely charged side chains are way above the cut-off distance (4.0 °A) as used in [34]. As such, the electrostatic interaction between B_ARG_36 and A_GLU_68 is not categorized as salt bridge [34] here.

**Table 3:** Interfacial residue pair distance analysis of <u>PDB entry: 4ZGM</u> [24]. In this table, the residue naming scheme is the same as Table 1, and the residue ID num-bering scheme is described in Table 1 of supplementary file **supps.pdf**, and chains A and B represent GLP-1R ECD and semaglutide, respectively.

| PDB ID | Residue A | Atom A | Residue B | Atom B | Distance (Å) |
| --- | --- | --- | --- | --- | --- |
| 4ZGM | B_ARG_36 | NH1 | A_GLU_68 | OE1 | 4.147 |
| 4ZGM | B_ARG_36 | NH1 | A_GLU_68 | OE2 | 4.835 |
| 4ZGM | B_ARG_36 | NH2 | A_GLU_68 | OE1 | 4.893 |
| 4ZGM | B_ARG_36 | NH2 | A_GLU_68 | OE2 | 6.269 |

### Rational design of semaglutide analogues with improved GLP-1R ECD binding affinity

In 2021, I introduced a manually designed Val-Arg exchange into the peptide backbone of semaglutide to improve its GLP-1R ECD binding affinity [27]. In this study, an automated high-throughput approach was employed for a systemic amino acid scanning of semaglutide’s backbone to enhance GLP-1R ECD affinity and GLP-1R activation thereafter. With the **Modigy** [28] workflow, a total of 564 amino acid sequences were computationally designed with three missense mutations according to the sequence template of semaglutide backbone in Figure 1, where Modeller [29] was used to generate at least 5000 homology complex structural models of GLP-1R ECD-semaglutide analogue with PDB entry 4ZGM [24, 35] as the structural template, and afterwards, for each structural model, Prodigy [30, 31] was used for the structural biophysics-based calculations of K_d_between semaglutide analogues and GLP-1R ECD.

For the 564 semaglutide analogues, supplementary file **sema.txt** provides a summary of:

1. an identification number of the backbone sequence, i.e., from 1 to 564;
2. three site-specific missense mutations introduced into the peptide backbone;
3. three site-specific missense mutations sorted according to their residue IDs;
4. original sequence of the peptide backbone of semaglutide, as in Figure 1;
5. mutant sequence of the peptide backbone of semaglutide;
6. an amino acid sequence alignment of the original and the mutant;
7. a statistical summary of the ligand-receptor binding affinity K_d_values between native semaglutide and GLP-1R ECD (PDB entry 4ZGM [24, 35]);
8. a statistical summary of the ligand-receptor binding affinity K_d_values between mutant semaglutide and GLP-1R ECD.

In supplementary file **sema.txt**, a statistical summary includes the average, the standard deviation, the maximum and the minimum of the ligand-receptor binding affinity K_d_values, where the average K_d_was used to rank the 564 semaglutide analogues, with the average K_d_of native semaglutide as the baseline. Of note, for three site-specific missense mutations in supplementary file **sema.txt**, the residue ID numbering scheme for semaglutide’s peptide backbone is from 1 to 28, as included in Table 1 of supplementary file **supps.pdf**, instead of from 7 to 37 as in Figure 1.

### Rational design of semaglutide analogues: a biophysical perspective

In supplementary file **sema.txt**, semaglutide mutant 524 is ranked the first due to its highest average GLP-1R ECD binding affinity (K_d_). Chemically, semaglutide mutant 524 is computationally designed as the peptide backbone of semaglutide (Figure 1) with three missense mutations, i.e., I20B Q, L23B R and V24B N (Table 1 of supplementary file **supps.pdf**). To further test if the three missense mutations do lead to a significantly increased GLP-1R ECD binding affinity, the **Modigy** [28] workflow was conducted for 10,000 times for both native semaglutide and semaglutide mutant 524, generating 10000 structural models built by Modeller [29] each for native semaglutide and semaglutide mutant 524. Afterwards, the 20,000 structural models were subject to ligand-receptor K_d_calculations by Prodigy [30, 31].

Of the two sets of K_d_data, a statistical analysis indicates a clear difference between them, where neither dataset follows a normal distribution (p-values *≈* 0). Given the non-normality of the data (Figure 2), a Wilcoxon test was used, yielding a final p-value of approximately 0, strongly suggesting that the two datasets are extremely different from each other. Taken together, as shown also in Figure 2, compared with that (3.278 *×* 10^−6^M) of native semaglutide (PDB entry 4ZGM [24]), semaglutide mutant 524 possesses a significantly higher (at least one degree of order) GLP-1R ECD binding affinity (1.462 *×* 10^−7^M) to enhance GLP-1R activation.

**Figure 2.**
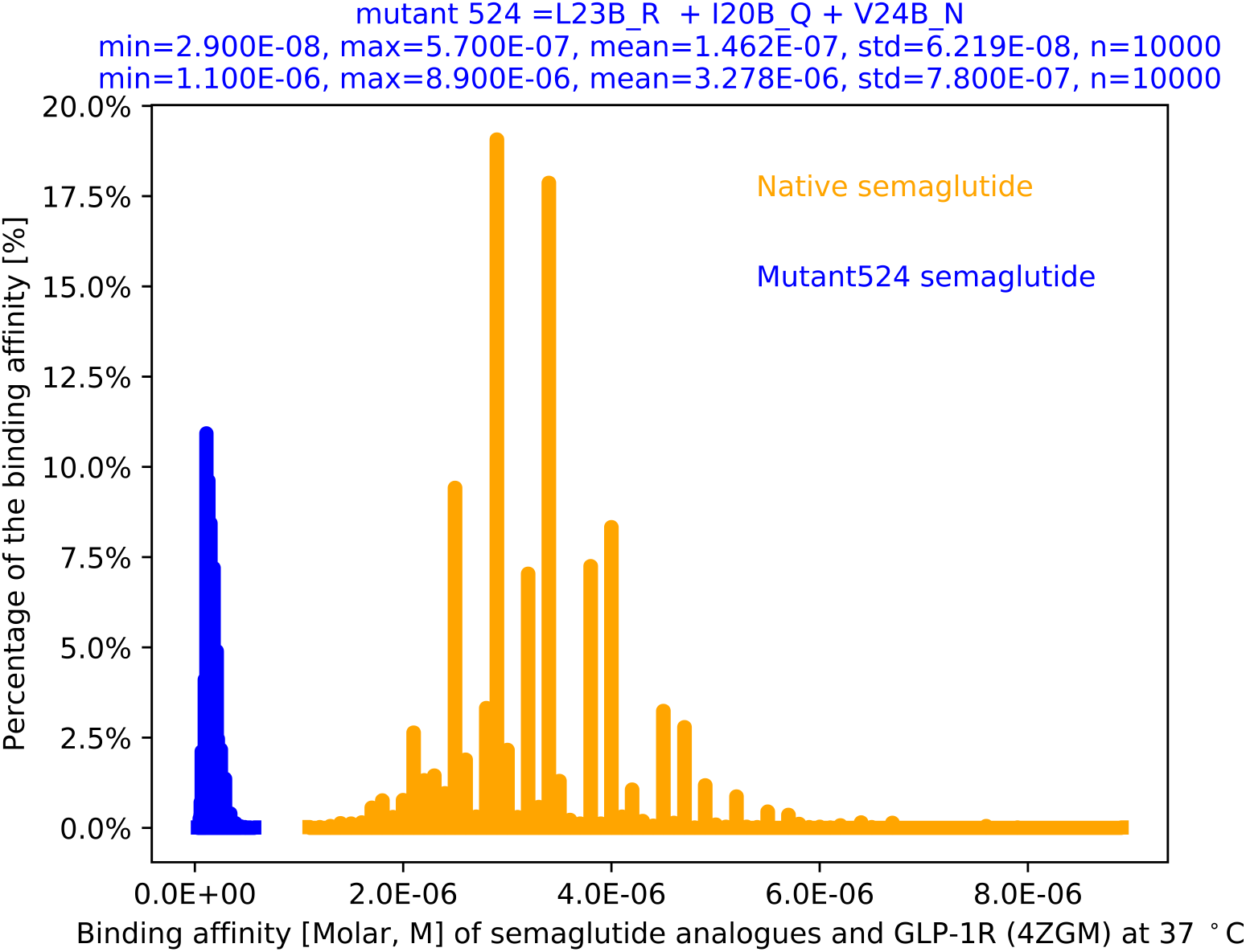
Distributions of the GLP-1R ECD binding affinities of native (the orange histogram) semaglutide (<u>PDB ID: 4ZGM</u> [24]) and semaglutide mutant 524 (the blue histogram). In the title (top) of this figure, L23B R, I20B Q and V24B R represent three positions where site-specific missense mutations are introduced to the peptide backbone of semaglutide to improve its GLP-1R ECD binding affinity.

To further investigate how the three missense mutations (I20B Q, L23B R and V24B N, Figure 2) led to a significantly increased GLP-1R ECD binding affinity, a comprehensive structural biophysical analysis [34] was conducted for the 10000 structural models of semaglutide mutant 524 built by Modeller [29]. As mentioned above, the role of charge-charge interactions is only as important as 4.76% (2/42, Table 4 of supplementary file **supps.pdf**) in the stabilization of the complex structure of GLP-1R ECD and semaglutide [24], including only two interfacial salt bridges (Table 1) and two interfacial hydrogen bonds (Table 2). Hence, the comprehensive structural biophysical analysis [34] here focuses primarily on the electrostatic interactions at the binding interface between GLP-1R ECD and semaglutide or its analogue (mutant 524).

**Table 4:** Interfacial salt bridge analysis [34] of the 4452^th^structural model (supplementary file **4452.pdb**) of semaglutide mutant 524, the first in supplementary file **sema.txt**. In this table, the residue naming scheme is the same as Table 1, and the residue ID numbering scheme is described in Table 1 of supplementary file **supps.pdf**, and chains A and B represent GLP-1R ECD and semaglutide, respectively, while the one row with yellow background represents the interfacial salt bridge also identified in PDB entry 4ZGM [24, 35].

| PDB file | Residue A | Atom A | Residue B | Atom B | Distance (Å) |
| --- | --- | --- | --- | --- | --- |
| 4452 | B_LYS_117 | NZ | A_GLU_100 | OE2 | 3.505 |
| 4452 | B_ARG_123 | NH1 | A_GLU_40 | OE1 | 3.417 |
| 4452 | B_ARG_127 | NH1 | A_GLU_40 | OE1 | 3.122 |
| 4452 | B_ARG_127 | NH1 | A_GLU_40 | OE2 | 4.023 |

Specifically, one structural model stood out, i.e., the 4452^th^structural model (supplementary file **4452.pdb**) of semaglutide mutant 524, where four salt bridges were identified at the binding interface of GLP-1R ECD and semaglutide mutant 524, as listed in Table 4. Among the four, three new interfacial salt bridges (Figures 3 and 4) were found to be formed due to the missense mutations introduced into semaglutide’s backbone (including in particular L23B R), whereas only one interfacial salt bridge (yellow row in Table 4) was found to be formed and the same as the salt bridge (Table 1) identified in PDB entry 4ZGM [24, 35].

**Figure 3.**
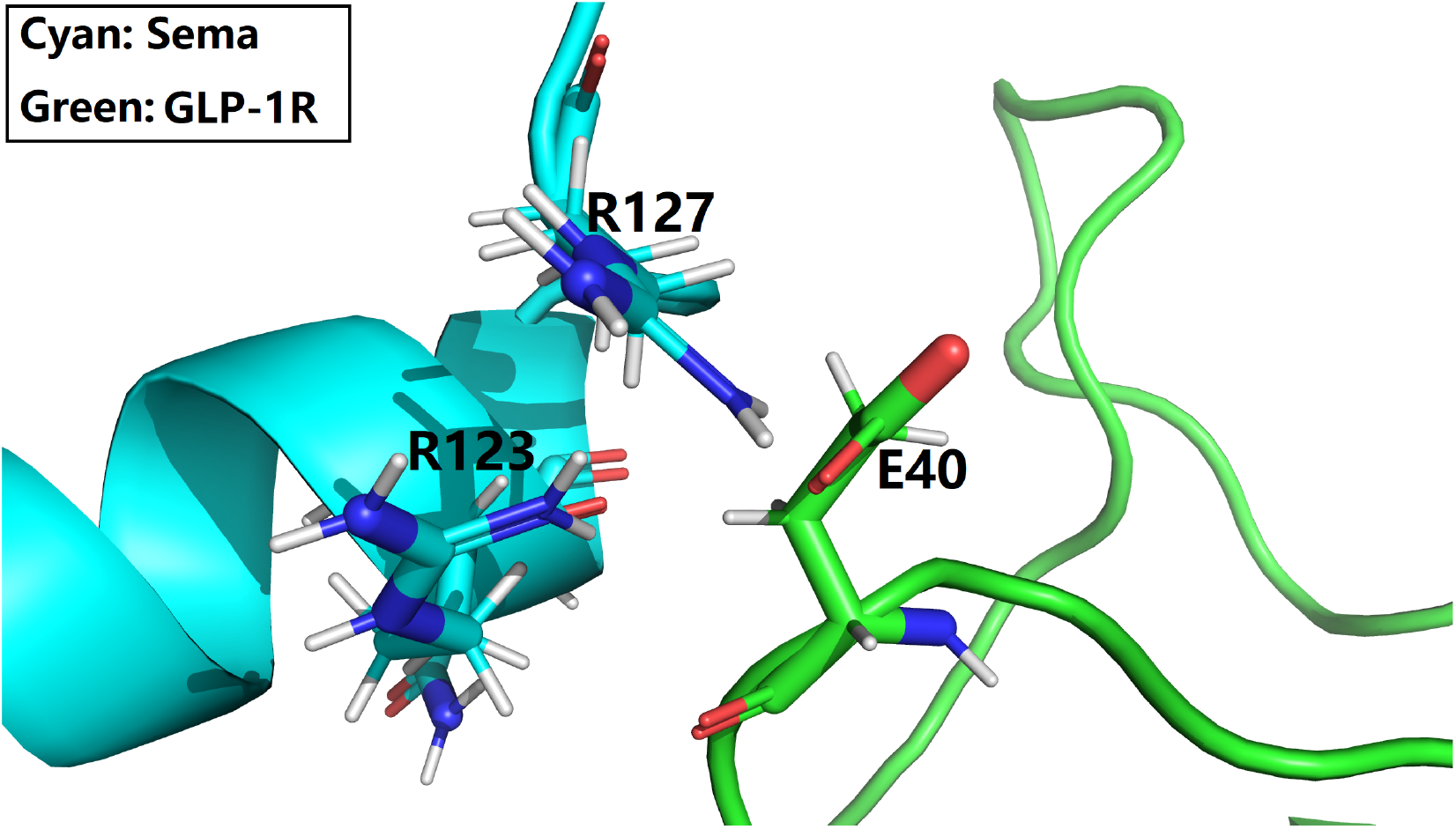
A zoomed-in view of the binding interface of GLP-1R and semaglutide mutant 4452 (Figure 2). This figure is prepared with PyMol [42] with supplementary file **4452.pdb**, and the details of the interfacial salt bridges are included in Table 4.

**Figure 4.**
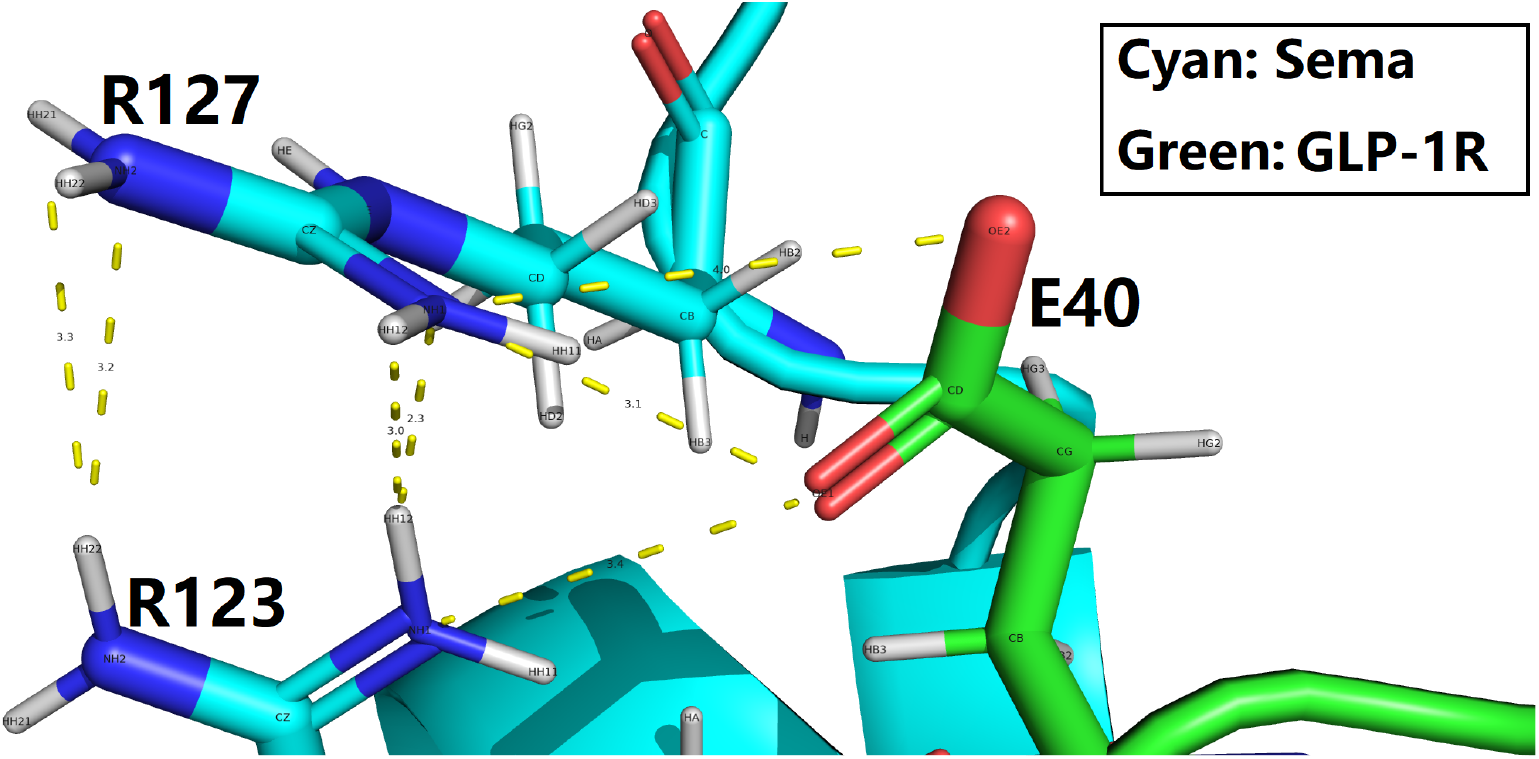
Salt bridges at the binding interface of GLP-1R and semaglutide mutant 4452 (Figure 2). This figure is prepared with PyMol [42] with supplementary file **4452.pdb**, the details of the interfacial salt bridges (yellow dotted lines) are included in Table 4, and the details of other H-bond-like electrostatic interactions are included in Table 5.

From Figures 3 and 4, it is quite clear that there is a cluster of three charged residues, i.e., Arg123 and Arg127 of semaglutide’s backbone, and Glu40 of GLP-1R ECD, sitting at the binding interface of the semaglutide-GLP-1R ECD complex structure. Here, both Arg123 and Arg127 of semaglutide’s backbone (cyan cartoon, Figures 3 and 4) formed interfacial salt bridges with Glu40 of GLP-1R ECD (green cartoon, Figures 3 and 4), the details of which are included in Table 4 and shown in Figure 4.

**Table 5:** Interfacial hydrogen bond analysis of the 4452^th^structural model (supplementary file **4452.pdb**) of semaglutide mutant 524. In this table, the residue naming scheme is the same as Table 1, and the residue ID numbering scheme is described in Table 1 of supplementary file **supps.pdf**, and chains A and B represent GLP-1R ECD and semaglutide, respectively, while ∠*ADH* represents the angle formed by acceptor (A), donor (D) and hydrogen (H).

| PDB | Acceptor (A) | Donor (D) | Hydrogen (H) | D-A (Å) | H-A (Å) | $\angle ADH(^{\circ})$ |
| --- | --- | --- | --- | --- | --- | --- |
| 4452 | NH1, B_ARG_127 | NH1, B_ARG_123 | HH12, B_ARG_123 | 3.23 | 2.29 | 16.94 |
| 4452 | NE, B_ARG_127 | NH1, B_ARG_123 | HH12, B_ARG_123 | 3.19 | 2.36 | 28.90 |

Here, of note, as described in Tables 2 and 3 of supplementary file **supps.pdf**,

1. B_LYS_117 of semaglutide mutant 524 (supplementary file **4452.pdb**) corresponds to B_LYS_26 of native semaglutide (PDB entry 4ZGM [24, 35]);
2. B_ARG_123 of semaglutide mutant 524 (supplementary file **4452.pdb**) corresponds to B_LEU_32 of native semaglutide (PDB entry 4ZGM [24, 35]);
3. B_ARG_127 of semaglutide mutant 524 (supplementary file **4452.pdb**) corresponds to B_ARG_36 of native semaglutide (PDB entry 4ZGM [24, 35]);
4. A_GLU_100 of semaglutide mutant 524 (supplementary file **4452.pdb**) corresponds to A_GLU_128 of native semaglutide (PDB entry 4ZGM [24, 35]);
5. A_GLU_40 of semaglutide mutant 524 (supplementary file **4452.pdb**) corresponds to A_GLU_68 of native semaglutide (PDB entry 4ZGM [24, 35]).

From Figures 3 and 4, in particular, it is obvious that the newly formed Arg123-Glu40 salt bridge is due to the missense mutation L23B R (Figure 2), i.e., the substitutin of the leucine with a positively charged arginine at position 32 (instead of 23, Table 1 of supplementary file **supps.pdf**) of semaglutide’s backbone (Figure 2). Moreover, the newly formed Arg127-Glu40 salt bridge is at least partly due to the newly formed Arg123-Glu40 salt bridge, i.e., the presence of a positively charged arginine at position 32 of semaglutide’s backbone (Figure 2), because in PDB entry 4ZGM [24, 35], the distances (Table 3) between the oppositely charged side chains of A_GLU_68 and B_ARG_36 are far beyond the cut-off distance (4.0 °A) for salt bridge analysis as used in [34]. While in the case of of semaglutide mutant 524 (supplementary file **4452.pdb**) here, one clear cut interfacial salt bridge was formed at 3.122 °A (Table 4) between Arg127 in semaglutide [24, 35]) and Glu40 of GLP-1R ECD, and another one also categorized here as an interfacial salt bridge at 4.023 °A, as the inter-actomic distance (4.023 °A, Table 4) equals almost the cut-off distance (4.0 °A) for salt bridge analysis as used in [34].

In Figures 3 and 4, the cluster of three charged residues consists of Arg123, Arg127 and Glu40, where the two arginines possess positively charged side chains, while the glutamate (Glu40) possesses a negatively charged side chain. Energetically, it is not favorable for Arg123 and Arg127 to be too close to each other due to charge-charge repulsion, despite the presence of a negatively charged Glu40 which attracts them. Of interest is the existence of two interfacial hydrogen bonds (Table 5) between the side chains of Arg123 and Arg127, which constitute an additional set of attractive forces for the structural stability of the cluster of three charged residues (Figures 3 and 4) consisting of Arg123, Arg127 and Glu40. As such, these two interfacial hydrogen bonds (Table 5), along with the three interfacial salt bridges (Figures 3 and 4) between Arg123, Arg127 and Glu40, led to the establishment of a delicate structural electrostatic balance, while maintaining the structural stabilization of the complex of GLP-R and semaglutide analogues, where such a delicate structural balance in hindsight entails a paradigm shift from the manual design approach in 2011 [27] to an automated systemic approach as described in this manuscript, as it does not appear so straightforwad for a manual design approach to uncover a such a delecate structural balance at a ligand-receptor binding interface [24].

Overall, from an molecular energetic point of view, the formation of these three new interfacial salt bridges (Figures 3 and 4) act like three electrostatic clips to further stabilize the ligand-receptor complex structure, and contribute favorably to the binding affinity of semaglutide mutant 542 to GLP-1R ECD, with the potential of the semaglutide analogue binding stronger and thereby leading to stronger GLP-1R activation [24]. Yet, it remains an open question of whether there is still room of optimization of the structural energetic (both electrostatics and hydrophobics) balance at the binding interface of GLP-1R ECD and semaglutide analogues, because afterall only three missense mutations were introduced into the peptide backbone of semaglutide with the Modigy [28] workflow here.

### An electrostatic scaffold for the design of newer GLP-1R agonists

As mentioned above, the role of charge-charge interactions is only as important as 4.76% (2/42, Table 4 of supplementary file **supps.pdf**) in the stabilization of the complex structure of GLP-1R ECD and semaglutide [24], including only two interfacial salt bridges (Table 1) and two interfacial hydrogen bonds (Table 2). As such, with 564 semaglutide analogues (supplementary file **sema.txt**) in place, a new set of Modigy [28] workflow was conducted with four missense mutations introduced into the peptide backbone of semaglutide, leading to the identfication of a structural electrostatic scaffold at the binding interface of GLP-1R ECD and semaglutide analogue A747 (supplementary file **A747.pdb**) with four site-specific missense mutations, namely L23B_R, I20B_Q, V24B_R and R25B_A.

Here, of note, this new Modigy [28] workflow with four missense mutations was only carried out for those key residue positions in semaglutide’s backbone and incomplete, because a complete Modigy [28] workflow here would require a total of 3,276,000,000 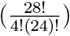 times of structural modeling by Modeller [29] and ligand-receptor binding affinity (K_d_) calculations by Prodigy [30, 31], where the size of the computational task is too huge even for a traditional HPC facility [28]. In addition, as described in Tables 1 and 3 of supplementary file **supps.pdf**,

1. I20B_Q represents a substitution of B_ILE_29 of native semaglutide (PDB entry 4ZGM [24, 35]) with B_GLN_120 in supplementary file **A747.pdb**;
2. L23B_R represents a substitution of B_LEU_32 of native semaglutide (PDB entry 4ZGM [24, 35]) with B_ARG_123 in supplementary file **A747.pdb**;
3. V24B_R represents a substitution of B_VAL_33 of native semaglutide (PDB entry 4ZGM [24, 35]) with B_ARG_124 in supplementary file **A747.pdb**;
4. R25B_A represents a substitution of B_ARG_34 of native semaglutide (PDB entry 4ZGM [24, 35]) with B_ALA_125 in supplementary file **A747.pdb**. Interestingly, this mutation along with V24B_R are equivalent to the manually designed Val-Arg exchange [] plus a Val-Ala substitution at position 34 native semaglutide (PDB entry 4ZGM [24, 35]).

With this partial Modigy [28] workflow, a total of 9 interfacial salt bridges (Table 6) were identified at the binding interface of GLP-1R ECD and semaglutide analogue A747 (supplementary file **A747.pdb**), in comparison with 2 interfacial salt bridges (Table 1) for PDB entry 4ZGM [24, 35]), and in comparison with 7 interfacial salt bridges (Table 4) for supplementary file **4452.pdb**. Specifically, three are a total of 7 newly formed interfacial salt bridges (Table 6) which are due to the four site-specific missense mutations L23B R, I20B Q, V24B R and R25B A introduced into the peptide backbone (Figure 1) of semaglutide.

**Table 6:**
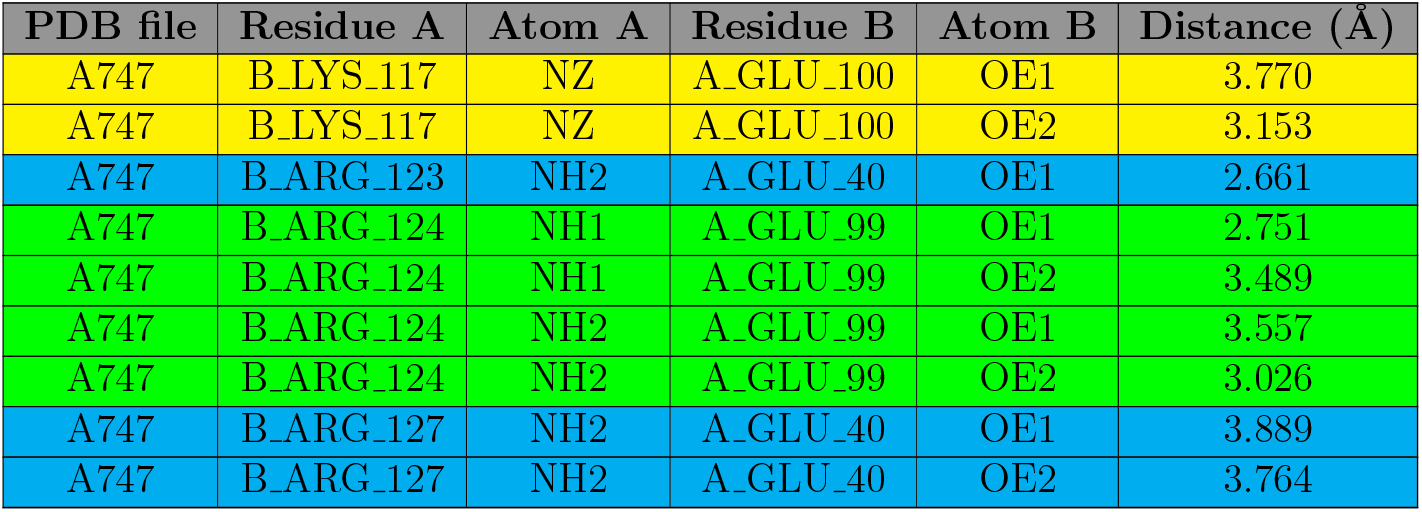
Interfacial salt bridging network analysis of semaglutide analogue A747 in complex with GLP-1R ECD (supplementary file **A747.pdb**. In this table, the residue naming scheme is the same as Table 1, and the residue ID numbering scheme is described in Table 1 of supplementary file **supps.pdf**, and chains A and B represent GLP-1R ECD and semaglutide, respectively, while rows with yellow, green and cyan backgrounds represent the first (1 in Figure 5), the second (2 in Figure 5) and the third (3 in Figure 5) sets of interfacial salt bridges, as shown in Figure 5 and also in the Graphical Abstract of this manuscript.

From a structural point of view, the nine interfacial salt bridges (Table 6) constitute a structural electrostatic scaffold (Figure 5) at the binding interface of GLP-1R ECD and semaglutide analogue A747, which consists of three sets of interfacial salt bridges, as designated by three red numbers (1, 2 and 3) within black circles in Figure 5, and also in Figures 1-5 in supplementary file **supps.pdf**. Moreover, the role of charge-charge interactions is as important as 18.18% (8/44, Table 5 in supplementary file **supps.pdf**) for semaglutide analogue A747 to GLP-1R ECD, compared with 4.76% (2/42, Table 4 in supplementary file **supps.pdf**) for native semaglutide to GLP-1R ECD, in the stabilization of the semaglutide backbone in complex with GLP-1R ECD (<u>PDB ID: 4ZGM</u> [24]).

**Figure 5.**
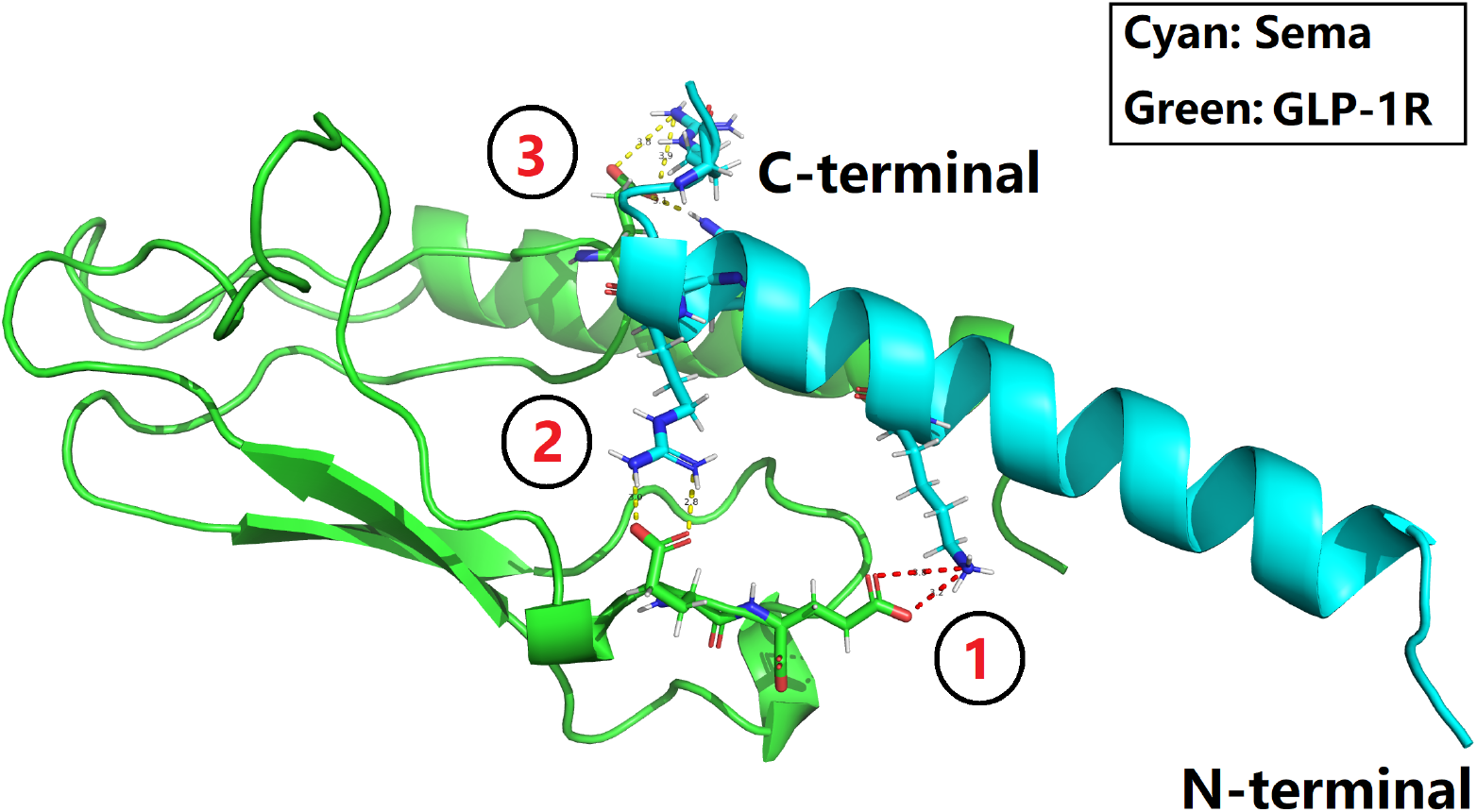
An overview of the complex structure of GLP-1R and semaglutide mutant A747. In this figure, the three red numbers (1, 2 and 3) within black circles represent the three sets of interfacial salt bridges (red and yellow dotted lines) of GLP-1R and semaglutide mutant A747, which constitute an electrostatic scaffold for the design of next-generation GLP-1R agonists with improved GLP-1R ECD affinity. This figure is prepared with PyMol [42] with supplementary file **A747.pdb**, with details of the three sets of interfacial salt bridges included in Table 6.

## Conclusion

In summary, this article reports a structural biophysics-driven computational framework, i.e., the **Modigy** workflow [28], for the design of semaglutide analogues (Figure 6), emphasizing structural and electrostatic optimization to enhance GLP-1R activation, to address the concern of the market with respect to the weight loss efficacy of CargiSema developed by Novo Nordisk. In addition to 564 semaglutide analogues (supplementary file **sema.txt**) with site-specific missense mutations, this study delves into the sequence space of GLP-1R ECD and semaglutide, and uncovers an electrostatic scaffold of intermolecular salt bridges at the ligand-receptor binding interface, which help stabilize the complex structure to enhance GLP-1R activation, providing a robust framework for the development of next-generation GLP-1R agonists with improved efficacy [43–46].

**Figure 6.**
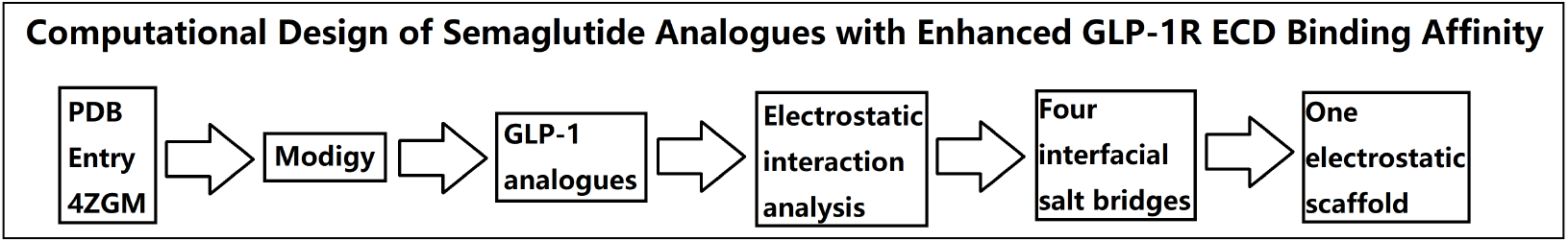
A flowchart overview of the material, methods, results, data analysis and conclusion of this study. In this figure, **Modigy** [28] is an abbreviation of Modeller [29] and Prodigy [30, 31].

## Discussion

Originally coined in 2024 [28], **Modigy** is defined as an abbreviation of Modeller [29] and Prodigy [30, 31] to represent an in silico high-throughput generation of structural and intermolecular binding affinity (K_d_) data [28]. Since its inception, the **Modigy** workflow aims to contribute to structure-based and biophysics-driven biomolecular discovery and design, making use of the experimental complex structures deposited in Protein Data Bank, e.g., ligand-receptor, antibody-antigen, etc. As such, for continued the development of next-generation GLP-1R agonists (e.g., CagriSema [20] and IcoSema [15]) with improved efficacy [43–46], the **Modigy** workflow is applicable not just for GLP-1R and GLP-1 (analogues), but also for amylin [47, 48], glucosedependent insulinotropic polypeptide receptor (GIPR) [49], glucagon receptor (GCGR) [50], and beyond, so long as an experimental complex structure is already deposited in the Protein Data Bank [36] for the direct interacting ligand and receptor [51].

Since the regulatory approval of semaglutide in 2017, the obesity and diabetes drug market has become highly competitive, led by Novo Nordisk and Eli Lilly. For instance, Novo Nordisk’s semaglutide-based drugs (Ozempic, Wegovy) face competition from Eli Lilly’s tirzepatide-based Mounjaro and Zepbound, which have demonstrated superior weight loss efficacy, while Novo Nordisk’s CagriSema has recently shown promising but not market-leading results, further intensifying industry competition. Both companies are expanding production and clinical trials to solidify their positions, as on December 6, 2024, the European Commission approved Novo Holdings’ acquisition of Catalent and related manufacturing sites for weight-loss drug production.

While pharmaceutical companies must balance marked competition and patient needs, longterm success depends on delivering continuously really effective, safe, and innovative treatments for patients. As such, given that the R&D process of a drug is itself a continued multi-paramater optimization process, this study here does not focus primarily on marked/product competition. Instead, this study here focuses primarily on continued exploration of the uncharted territory, which is the unexplored sequence/chemical space of GLP-1R agonists, for the development of next-generation GLP-1R agonists with improved efficacy [43–46], safety and hopefully cost, too.

### Declaration of generative AI and AI-assisted technologies in the writing process

During the preparation of this work, the author used OpenAI’s ChatGPT for the choice of words and for the organization of a few sentences, in order to improve the readability of the manuscript. After using this tool, the author reviewed and edited the content as needed and takes full responsibility for this manuscript.

## Supporting information

supplementary tables, figures and PDB files of the computationally designed semaglutide analogues.

## References

1. Sims EK, Carr ALJ, Oram RA, DiMeglio LA, Evans-Molina C. 100 years of insulin: celebrating the past, present and future of diabetes therapy. Nature Medicine. 2021;27(7):1154–1164. doi:10.1038/s41591-021-01418-2.

2. Vecchio I, Tornali C, Bragazzi NL, Martini M. The Discovery of Insulin: An Important Milestone in the History of Medicine. Frontiers in Endocrinology. 2018;9. doi:10.3389/fendo.2018.00613.

3. Hirsch IB. Insulin Analogues. New England Journal of Medicine. 2005;352(2):174–183.

4. Scapin G, Dandey VP, Zhang Z, Strickland C, Potter CS, Carragher B. Insulin Receptor ectodomain in complex with one insulin molecule; 2018.

5. Li W. Delving Deep into the Structural Aspects of the BPro28-BLys29 Exchange in Insulin Lispro: A Structural Biophysical Lesson. 2020;.

6. Knudsen LB, Lau J. The Discovery and Development of Liraglutide and Semaglutide. Frontiers in Endocrinology. 2019;10:1–32.

7. Bucheit JD, Pamulapati LG, Carter N, Malloy K, Dixon DL, Sisson EM. Oral Semaglutide: A Review of the First Oral Glucagon-Like Peptide 1 Receptor Agonist. Diabetes Technology & Therapeutics. 2020;22(1):10–18.

8. Rodbard HW, Rosenstock J, Canani LH, Deerochanawong C, Gumprecht J, Lindberg SØ, et al. Oral Semaglutide Versus Empagliflozin in Patients With Type 2 Diabetes Uncontrolled on Metformin: The PIONEER 2 Trial. Diabetes Care. 2019;42(12):2272–2281.

9. Xiao Y, Sun L. Semaglutide in weight management. The Lancet. 2019;394(10205):1226.

10. Anderson SL, Beutel TR, Trujillo JM. Oral semaglutide in type 2 diabetes. Journal of Diabetes and its Complications. 2020;34(4):107520.

11. Røder ME. Clinical potential of treatment with semaglutide in type 2 diabetes patients. Drugs in Context. 2019;8:1–11.

12. Aroda VR, Rosenstock J, Terauchi Y, Altuntas Y, Lalic NM, Villegas ECM, et al. PIONEER 1: Randomized Clinical Trial of the Efficacy and Safety of Oral Semaglutide Monotherapy in Comparison With Placebo in Patients With Type 2 Diabetes. Diabetes Care. 2019;42(9):1724–1732.

13. Pratley R, Amod A, Hoff ST, Kadowaki T, Lingvay I, Nauck M, et al. Oral semaglutide versus subcutaneous liraglutide and placebo in type 2 diabetes (PIONEER 4): a randomised, double-blind, phase 3a trial. The Lancet. 2019;394(10192):39–50.

14. Husain M, Birkenfeld AL, Donsmark M, Dungan K, Eliaschewitz FG, Franco DR, et al. Oral Semaglutide and Cardiovascular Outcomes in Patients with Type 2 Diabetes. New England Journal of Medicine. 2019;381(9):841–851.

15. Kalra S, Bhattacharya S, Kapoor N. Contemporary Classification of Glucagon-Like Peptide 1 Receptor Agonists (GLP1RAs). Diabetes Therapy. 2021;12(8):2133–2147.

16. Seetharaman R. IcoSema’s leap forward: new data from COMBINE 3 paves the way. J Basic Clin Physiol Pharmacol. 2024;.

17. Westergaard L, Alifrangis L, Buckley ST, Coester HV, Klitgaard T, Kristensen NR, et al. Pharmacokinetic properties of a once-weekly fixed-ratio combination of insulin icodec and semaglutide compared with separate administration of each component in individuals with type 2 diabetes mellitus. Clin Drug Investig. 2024;44(11):849–861.

18. Lingvay I, Asong M, Desouza C, Gourdy P, Kar S, Vianna A, et al. Once-weekly insulin icodec vs once-daily insulin degludec in adults with insulin-naive type 2 diabetes: The ONWARDS 3 randomized clinical trial. JAMA. 2023;330(3):228–237.

19. Philis-Tsimikas A, Asong M, Franek E, Jia T, Rosenstock J, Stachlewska K, et al. Switching to once-weekly insulin icodec versus once-daily insulin degludec in individuals with basal insulin-treated type 2 diabetes (ONWARDS 2): a phase 3a, randomised, open label, multicentre, treat-to-target trial. Lancet Diabetes Endocrinol. 2023;11(6):414–425.

20. Apovian CM, McDonnell ME. CagriSema and the link between obesity and type 2 diabetes. The Lancet. 2023;402(10403):671–673. doi:10.1016/s0140-6736(23)01291-6.

21. Longwell CK, Hanna S, Hartrampf N, Sperberg RAP, Huang PS, Pentelute BL, et al. Identification of N-Terminally Diversified GLP-1R Agonists Using Saturation Mutagenesis and Chemical Design. ACS Chemical Biology. 2020;16(1):58–66. doi:10.1021/acschembio.0c00722.

22. Zhang H, Wu T, Wu Y, Peng Y, Wei X, Lu T, et al. Binding sites and design strategies for small molecule GLP-1R agonists. European Journal of Medicinal Chemistry. 2024;275:116632. doi:10.1016/j.ejmech.2024.116632.

23. Zhang X, Belousoff MJ, Liang YL, Danev R, Sexton PM, Wootten D. Structure and dynamics of semaglutide- and taspoglutide-bound GLP-1R-Gs complexes. Cell Reports. 2021;36(2):109374. doi:10.1016/j.celrep.2021.109374.

24. Lau J, Bloch P, Schäffer L, Pettersson I, Spetzler J, Kofoed J, et al. Discovery of the Once-Weekly Glucagon-Like Peptide-1 (GLP-1) Analogue Semaglutide. Journal of Medicinal Chemistry. 2015;58(18):7370–7380.

25. Dang T, Yu J, Cao Z, Zhang B, Li S, Xin Y, et al. Endogenous cell membrane interactome mapping for the GLP-1 receptor in different cell types. Nature Chemical Biology. 2024;doi:10.1038/s41589-024-01714-1.

26. Wright SC, Motso A, Koutsilieri S, Beusch CM, Sabatier P, Berghella A, et al. GLP-1R signaling neighborhoods associate with the susceptibility to adverse drug reactions of incretin mimetics. Nature Communications. 2023;14(1). doi:10.1038/s41467-023-41893-4.

27. Li W. Strengthening Semaglutide-GLP-1R Binding Affinity via a Val27-Arg28 Exchange in the Peptide Backbone of Semaglutide: A Computational Structural Approach. Journal of Computational Biophysics and Chemistry. 2021;20(05):495–499.

28. Li W. In Silico Generation of Structural and Intermolecular Binding Affinity Data with Reasonable Accuracy: Expanding Horizons in Drug Discovery and Design. 2024;doi:10.20944/preprints202405.1739.v1.

29. Webb B, Sali A. Protein Structure Modeling with MODELLER. In: Methods in Molecular Biology. Springer US; 2020. p. 239–255.

30. Vangone A, Bonvin AM. Contacts-based prediction of binding affinity in protein-protein complexes. eLife. 2015;4.

31. Xue LC, Rodrigues JP, Kastritis PL, Bonvin AM, Vangone A. PRODIGY: a web server for predicting the binding affinity of protein-protein complexes. Bioinformatics. 2016; p. btw514.

32. Gilbert MP, Pratley RE. GLP-1 Analogs and DPP-4 Inhibitors in Type 2 Diabetes Therapy: Review of Head-to-Head Clinical Trials. Frontiers in Endocrinology. 2020;11. doi:10.3389/fendo.2020.00178.

33. Deacon CF. Dipeptidyl peptidase 4 inhibitors in the treatment of type 2 diabetes mellitus. Nature Reviews Endocrinology. 2020;16(11):642–653. doi:10.1038/s41574-020-0399-8.

34. Li W. How do SMA-linked mutations of SMN1 lead to structural/functional deficiency of the SMA protein? PLOS ONE. 2017;12(6):e0178519.

35. Reedtz-Runge S. Crystal structure of Semaglutide peptide backbone in complex with the GLP-1 receptor extracellular domain; 2015.

36. Berman H, Henrick K, Nakamura H. Announcing the worldwide Protein Data Bank. Nature Structural & Molecular Biology. 2003;10(12):980–980.

37. Wu D, Sun J, Xu T, Wang S, Li G, Li Y, et al. Stacking and energetic contribution of aromatic islands at the binding interface of antibody proteins. Immunome Research. 2010;6(Suppl 1):S1.

38. Davis FP, Sali A. PIBASE: a comprehensive database of structurally defined protein interfaces. Bioinformatics. 2005;21(9):1901–1907.

39. Jubb HC, Pandurangan AP, Turner MA, Ochoa-Montaño B, Blundell TL, Ascher DB. Mutations at protein-protein interfaces: Small changes over big surfaces have large impacts on human health. Progress in Biophysics and Molecular Biology. 2017;128:3–13.

40. Wells JA, McClendon CL. Reaching for high-hanging fruit in drug discovery at protein-protein interfaces. Nature. 2007;450(7172):1001–1009.

41. Antolíková E, Žáková L, Turkenburg JP, Watson CJ, Hančlová I, Šanda M, et al. Non-equivalent Role of Inter- and Intramolecular Hydrogen Bonds in the Insulin Dimer Interface. Journal of Biological Chemistry. 2011;286(42):36968–36977.

42. DeLano WL. Pymol: An open-source molecular graphics tool. CCP4 Newsletter On Protein Crystallography. 2002;40:82–92.

43. Francis D, Chacko AM, Anoop A, Nadimuthu S, Venugopal V. In: Evolution of biosynthetic human insulin and its analogues for diabetes management. Elsevier; 2024. p. 191–256.

44. Rosenstock J, Juneja R, Beals JM, Moyers JS, Ilag L, McCrimmon RJ. The Basis for Weekly Insulin Therapy: Evolving Evidence With Insulin Icodec and Insulin Efsitora Alfa. Endocrine Reviews. 2024;45(3):379–413. doi:10.1210/endrev/bnad037.

45. Derewenda U, Derewenda Z, Dodson GG, Hubbard RE, Korber F. Molecular Structure of insulin: The insulin monomer and its assembly. British Medical Bulletin. 1989;45(1):4–18.

46. Slieker LJ, Brooke GS, DiMarchi RD, Flora DB, Green LK, Hoffmann JA, et al. Modifications in the B10 and B26-30 regions of the B chain of human insulin alter affinity for the human IGF-I receptor more than for the insulin receptor. Diabetologia. 1997;40(14):S54– S61.

47. Kruse T, Hansen JL, Dahl K, Schäffer L, Sensfuss U, Poulsen C, et al. Development of Cagrilintide, a Long-Acting Amylin Analogue. Journal of Medicinal Chemistry. 2021;64(15):11183–11194. doi:10.1021/acs.jmedchem.1c00565.

48. Li W. Identification of Electrostatic Hotspots at the Binding Interface of Amylin and Insulin-Degrading Enzyme: A Structural and Biophysical Investigation. 2024;doi:10.20944/preprints202402.0063.v1.

49. Li W, Zhou Q, Cong Z, Yuan Q, Li W, Zhao F, et al. Structural insights into the triple agonism at GLP-1R, GIPR and GCGR manifested by retatrutide. Cell Discovery. 2024;10(1). doi:10.1038/s41421-024-00700-0.

50. Rosenstock J, Bain SC, Gowda A, Jódar E, Liang B, Lingvay I, et al. Weekly Icodec versus Daily Glargine U100 in Type 2 Diabetes without Previous Insulin. New England Journal of Medicine. 2023;389(4):297–308.

51. Li W, Vottevor G. Towards a Truly General Intermolecular Binding Affinity Calculator for Drug Discovery & Design. 2023;doi:10.20944/preprints202208.0213.v2.

