## supplementary tables, figures and PDB files of the computationally designed semaglutide analogues. for "Structural Biophysics-Guided Computational Design of Semaglutide Analogues to Enhance GLP-1R Activation": grafic.pdf

March 27, 2025

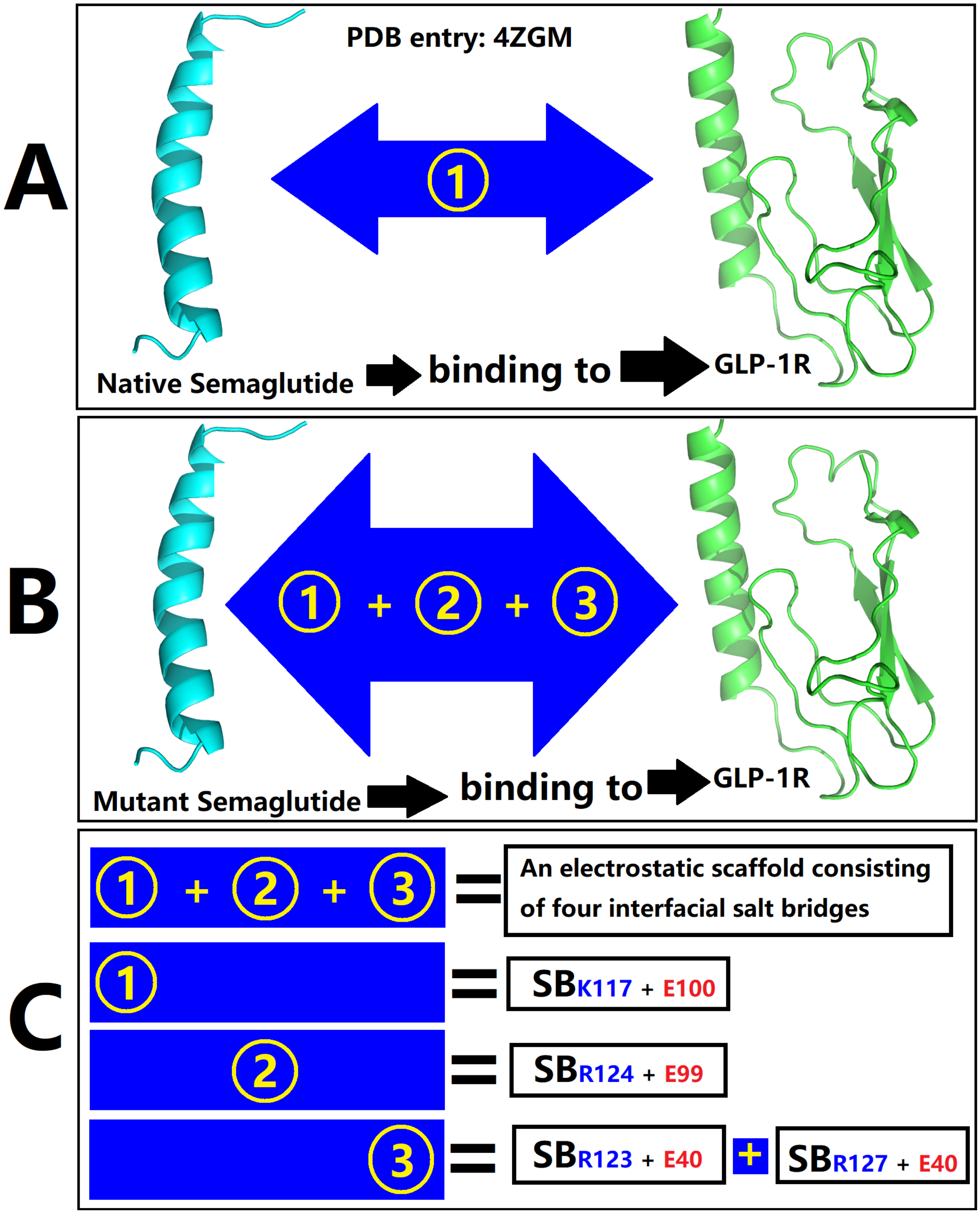


Figure 1: Structural biophysics-aided computational design of semaglutide analogues with improved GLP-1R extracellular domain (ECD) binding affinity through site-specific missense mutations introduced into the peptide backbone of semaglutide. In subfigures A and B, the cyan and the green cartoons are prepared with PyMol [1] with PDB entry 4GZM [2, 3], representing semaglutide and GLP-1R ECD, respectively. In subfigure C, numbers 1, 2 and 3 (yellow texts) represent three sets of salt bridges at the binding interface of GLP-1R ECD and semaglutide analogues, where positively and negatively charged residues are specified with blue and red texts in subfigure C, respectively, and **SB** is an abbreviation of salt bridge.

### References


[1] DeLano WL. Pymol: An open-source molecular graphics tool. CCP4 Newsletter On Protein Crystallography. 2002;40:82–92.

[2] Lau J, Bloch P, Schäffer L, Pettersson I, Spetzler J, Kofoed J, et al. Discovery of the Once-Weekly Glucagon-Like Peptide-1 (GLP-1) Analogue Semaglutide. Journal of Medicinal Chemistry. 2015;58(18):7370–7380.

[3] Reedtz-Runge S. Crystal structure of Semaglutide peptide backbone in complex with the GLP-1 receptor extracellular domain; 2015.
