## supplementary tables, figures and PDB files of the computationally designed semaglutide analogues. for "Structural Biophysics-Guided Computational Design of Semaglutide Analogues to Enhance GLP-1R Activation": supps.pdf

March 27, 2025

| ResName | ResID | Chain ID | ResName | ResID | Chain ID |
|---|---|---|---|---|---|
| 4ZGM | 4ZGM | 4ZGM | Modigy | Modigy | Modigy |
| GLY | 10 | B | GLY | 1 | B |
| THR | 11 | B | THR | 2 | B |
| PHE | 12 | B | PHE | 3 | B |
| THR | 13 | B | THR | 4 | B |
| SER | 14 | B | SER | 5 | B |
| ASP | 15 | B | ASP | 6 | B |
| VAL | 16 | B | VAL | 7 | B |
| SER | 17 | B | SER | 8 | B |
| SER | 18 | B | SER | 9 | B |
| TYR | 19 | B | TYR | 10 | B |
| LEU | 20 | B | LEU | 11 | B |
| GLU | 21 | B | GLU | 12 | B |
| GLY | 22 | B | GLY | 13 | B |
| GLN | 23 | B | GLN | 14 | B |
| ALA | 24 | B | ALA | 15 | B |
| ALA | 25 | B | ALA | 16 | B |
| LYS | 26 | B | LYS | 17 | B |
| GLU | 27 | B | GLU | 18 | B |
| PHE | 28 | B | PHE | 19 | B |
| ILE | 29 | B | ILE | 20 | B |
| ALA | 30 | B | ALA | 21 | B |
| TRP | 31 | B | TRP | 22 | B |
| LEU | 32 | B | LEU | 23 | B |
| VAL | 33 | B | VAL | 24 | B |
| ARG | 34 | B | ARG | 25 | B |
| GLY | 35 | B | GLY | 26 | B |
| ARG | 36 | B | ARG | 27 | B |
| GLY | 37 | B | GLY | 28 | B |

Table 1: Two amino acid residue ID numbering schemes of the peptide backbone of semaglutide in PDB entry: 4ZGM and in the **Modigy** [1] workflow. In this table, **ResName** represents the three-letter name of the amino acid residue, **ResID** represents the identification number of the amino acid residues in PDB entry: 4ZGM and in the **Modigy** [1] workflow, while **ChainID** specifies the molecular entity in PDB entry: 4ZGM and in the **Modigy** [1] workflow, with **A** for GLP-1R ECD and **B** for semaglutide. Here, the **Modigy** [1] workflow is defined as an abbreviation of Modeller [2] and Prodigy [3, 4] to represent an in silico high-throughput generation of structural and intermolecular binding affinity ($K_d$) data [1]. The three rows that are shaded in yellow represent three positions where site-specific missense mutations are introduced to the peptide backbone of semaglutide to improve its GLP-1R ECD binding affinity.

| ResName | ResID | Chain ID | ResName | ResID | Chain ID | ResName | ResID | Chain ID |
|---|---|---|---|---|---|---|---|---|
| 4ZGM | 4ZGM | 4ZGM | 4452 | 4452 | 4452 | A747 | A747 | A747 |
| GLY | 10 | B | GLY | 101 | B | GLY | 101 | B |
| THR | 11 | B | THR | 102 | B | THR | 102 | B |
| PHE | 12 | B | PHE | 103 | B | PHE | 103 | B |
| THR | 13 | B | THR | 104 | B | THR | 104 | B |
| SER | 14 | B | SER | 105 | B | SER | 105 | B |
| ASP | 15 | B | ASP | 106 | B | ASP | 106 | B |
| VAL | 16 | B | VAL | 107 | B | VAL | 107 | B |
| SER | 17 | B | SER | 108 | B | SER | 108 | B |
| SER | 18 | B | SER | 109 | B | SER | 109 | B |
| TYR | 19 | B | TYR | 110 | B | TYR | 110 | B |
| LEU | 20 | B | LEU | 111 | B | LEU | 111 | B |
| GLU | 21 | B | GLU | 112 | B | GLU | 112 | B |
| GLY | 22 | B | GLY | 113 | B | GLY | 113 | B |
| GLN | 23 | B | GLN | 114 | B | GLN | 114 | B |
| ALA | 24 | B | ALA | 115 | B | ALA | 115 | B |
| ALA | 25 | B | ALA | 116 | B | ALA | 116 | B |
| LYS | 26 | B | LYS | 117 | B | LYS | 117 | B |
| GLU | 27 | B | GLU | 118 | B | GLU | 118 | B |
| PHE | 28 | B | PHE | 119 | B | PHE | 119 | B |
| ILE | 29 | B | GLN | 120 | B | GLN | 120 | B |
| ALA | 30 | B | ALA | 121 | B | ALA | 121 | B |
| TRP | 31 | B | TRP | 122 | B | TRP | 122 | B |
| LEU | 32 | B | ARG | 123 | B | ARG | 123 | B |
| VAL | 33 | B | ASN | 124 | B | ARG | 124 | B |
| ARG | 34 | B | ARG | 125 | B | ALA | 125 | B |
| GLY | 35 | B | GLY | 126 | B | GLY | 126 | B |
| ARG | 36 | B | ARG | 127 | B | ARG | 127 | B |
| GLY | 37 | B | GLY | 128 | B | GLY | 128 | B |

Table 2: A vertical alignment of amino acid residues of the peptide backbone of semaglutide in PDB entry: 4ZGM, PDB file 4452.pdb and PDB file A747.pdb. In this table, **ResName** represents the three-letter name of the amino acid residue, **ResID** represents the identification number of the amino acid residues in PDB entry: 4ZGM, PDB file 4452.pdb and PDB file A747.pdb, while **ChainID** specifies the molecular entity in PDB entry: 4ZGM, PDB file 4452.pdb and PDB file A747.pdb, with **A** for GLP-1R ECD and **B** for semaglutide. The rows that are shaded in yellow represent three positions where site-specific missense mutations are introduced to the peptide backbone of semaglutide to improve its GLP-1R ECD binding affinity.

| ResName | ResID | Chain ID | ResName | ResID | Chain ID | ResName | ResID | Chain ID |
|---|---|---|---|---|---|---|---|---|
| 4ZGM | 4ZGM | 4ZGM | 4452 | 4452 | 4452 | A747 | A747 | A747 |
| THR | 29 | A | THR | 1 | A | THR | 1 | A |
| VAL | 30 | A | VAL | 2 | A | VAL | 2 | A |
| SER | 31 | A | SER | 3 | A | SER | 3 | A |
| LEU | 32 | A | LEU | 4 | A | LEU | 4 | A |
| TRP | 33 | A | TRP | 5 | A | TRP | 5 | A |
| GLU | 34 | A | GLU | 6 | A | GLU | 6 | A |
| THR | 35 | A | THR | 7 | A | THR | 7 | A |
| VAL | 36 | A | VAL | 8 | A | VAL | 8 | A |
| GLN | 37 | A | GLN | 9 | A | GLN | 9 | A |
| LYS | 38 | A | LYS | 10 | A | LYS | 10 | A |
| TRP | 39 | A | TRP | 11 | A | TRP | 11 | A |
| ARG | 40 | A | ARG | 12 | A | ARG | 12 | A |
| GLU | 41 | A | GLU | 13 | A | GLU | 13 | A |
| TYR | 42 | A | TYR | 14 | A | TYR | 14 | A |
| ARG | 43 | A | ARG | 15 | A | ARG | 15 | A |
| ARG | 44 | A | ARG | 16 | A | ARG | 16 | A |
| GLN | 45 | A | GLN | 17 | A | GLN | 17 | A |
| CYS | 46 | A | CYS | 18 | A | CYS | 18 | A |
| GLN | 47 | A | GLN | 19 | A | GLN | 19 | A |
| ARG | 48 | A | ARG | 20 | A | ARG | 20 | A |
| SER | 49 | A | SER | 21 | A | SER | 21 | A |
| LEU | 50 | A | LEU | 22 | A | LEU | 22 | A |
| THR | 51 | A | THR | 23 | A | THR | 23 | A |
| GLU | 52 | A | GLU | 24 | A | GLU | 24 | A |
| ASP | 53 | A | ASP | 25 | A | ASP | 25 | A |
| PRO | 54 | A | PRO | 26 | A | PRO | 26 | A |
| PRO | 55 | A | PRO | 27 | A | PRO | 27 | A |
| PRO | 56 | A | PRO | 28 | A | PRO | 28 | A |
| ALA | 57 | A | ALA | 29 | A | ALA | 29 | A |
| THR | 58 | A | THR | 30 | A | THR | 30 | A |
| ASP | 59 | A | ASP | 31 | A | ASP | 31 | A |
| LEU | 60 | A | LEU | 32 | A | LEU | 32 | A |
| PHE | 61 | A | PHE | 33 | A | PHE | 33 | A |
| CYS | 62 | A | CYS | 34 | A | CYS | 34 | A |
| ASN | 63 | A | ASN | 35 | A | ASN | 35 | A |
| ARG | 64 | A | ARG | 36 | A | ARG | 36 | A |
| THR | 65 | A | THR | 37 | A | THR | 37 | A |
| PHE | 66 | A | PHE | 38 | A | PHE | 38 | A |
| ASP | 67 | A | ASP | 39 | A | ASP | 39 | A |
| GLU | 68 | A | GLU | 40 | A | GLU | 40 | A |
| TYR | 69 | A | TYR | 41 | A | TYR | 41 | A |
| ALA | 70 | A | ALA | 42 | A | ALA | 42 | A |
| CYS | 71 | A | CYS | 43 | A | CYS | 43 | A |
| TRP | 72 | A | TRP | 44 | A | TRP | 44 | A |
| PRO | 73 | A | PRO | 45 | A | PRO | 45 | A |
| ASP | 74 | A | ASP | 46 | A | ASP | 46 | A |
| GLY | 75 | A | GLY | 47 | A | GLY | 47 | A |
| GLU | 76 | A | GLU | 48 | A | GLU | 48 | A |
| PRO | 77 | A | PRO | 49 | A | PRO | 49 | A |
| GLY | 78 | A | GLY | 50 | A | GLY | 50 | A |
| SER | 79 | A | SER | 51 | A | SER | 51 | A |
| PHE | 80 | A | PHE | 52 | A | PHE | 52 | A |
| VAL | 81 | A | VAL | 53 | A | VAL | 53 | A |
| ASN | 82 | A | ASN | 54 | A | ASN | 54 | A |
| VAL | 83 | A | VAL | 55 | A | VAL | 55 | A |
| SER | 84 | A | SER | 56 | A | SER | 56 | A |
| CYS | 85 | A | CYS | 57 | A | CYS | 57 | A |
| PRO | 86 | A | PRO | 58 | A | PRO | 58 | A |
| TRP | 87 | A | TRP | 59 | A | TRP | 59 | A |
| TYR | 88 | A | TYR | 60 | A | TYR | 60 | A |
| LEU | 89 | A | LEU | 61 | A | LEU | 61 | A |
| PRO | 90 | A | PRO | 62 | A | PRO | 62 | A |
| TRP | 91 | A | TRP | 63 | A | TRP | 63 | A |
| ALA | 92 | A | ALA | 64 | A | ALA | 64 | A |
| SER | 93 | A | SER | 65 | A | SER | 65 | A |
| SER | 94 | A | SER | 66 | A | SER | 66 | A |
| VAL | 95 | A | VAL | 67 | A | VAL | 67 | A |
| PRO | 96 | A | PRO | 68 | A | PRO | 68 | A |
| GLN | 97 | A | GLN | 69 | A | GLN | 69 | A |
| GLY | 98 | A | GLY | 70 | A | GLY | 70 | A |

| | | | | | | | | |
|---|---|---|---|---|---|---|---|---|
| HIS | 99 | A | HIS | 71 | A | HIS | 71 | A |
| VAL | 100 | A | VAL | 72 | A | VAL | 72 | A |
| TYR | 101 | A | TYR | 73 | A | TYR | 73 | A |
| ARG | 102 | A | ARG | 74 | A | ARG | 74 | A |
| PHE | 103 | A | PHE | 75 | A | PHE | 75 | A |
| CYS | 104 | A | CYS | 76 | A | CYS | 76 | A |
| THR | 105 | A | THR | 77 | A | THR | 77 | A |
| ALA | 106 | A | ALA | 78 | A | ALA | 78 | A |
| GLU | 107 | A | GLU | 79 | A | GLU | 79 | A |
| GLY | 108 | A | GLY | 80 | A | GLY | 80 | A |
| LEU | 109 | A | LEU | 81 | A | LEU | 81 | A |
| TRP | 110 | A | TRP | 82 | A | TRP | 82 | A |
| LEU | 111 | A | LEU | 83 | A | LEU | 83 | A |
| GLN | 112 | A | GLN | 84 | A | GLN | 84 | A |
| LYS | 113 | A | LYS | 85 | A | LYS | 85 | A |
| ASP | 114 | A | ASP | 86 | A | ASP | 86 | A |
| ASN | 115 | A | ASN | 87 | A | ASN | 87 | A |
| SER | 116 | A | SER | 88 | A | SER | 88 | A |
| SER | 117 | A | SER | 89 | A | SER | 89 | A |
| LEU | 118 | A | LEU | 90 | A | LEU | 90 | A |
| PRO | 119 | A | PRO | 91 | A | PRO | 91 | A |
| TRP | 120 | A | TRP | 92 | A | TRP | 92 | A |
| ARG | 121 | A | ARG | 93 | A | ARG | 93 | A |
| ASP | 122 | A | ASP | 94 | A | ASP | 94 | A |
| LEU | 123 | A | LEU | 95 | A | LEU | 95 | A |
| SER | 124 | A | SER | 96 | A | SER | 96 | A |
| GLU | 125 | A | GLU | 97 | A | GLU | 97 | A |
| CYS | 126 | A | CYS | 98 | A | CYS | 98 | A |
| GLU | 127 | A | GLU | 99 | A | GLU | 99 | A |
| GLU | 128 | A | GLU | 100 | A | GLU | 100 | A |

Table 3: A vertical alignment of amino acid residues of GLP-1R ECD in PDB entry: 4ZGM, PDB file 4452.pdb and PDB file A747.pdb. In this table, **ResName** represents the three-letter name of the amino acid residue, **ResID** represents the identification number of the amino acid residues in PDB entry: 4ZGM, PDB file 4452.pdb and PDB file A747.pdb, while **ChainID** specifies the molecular entity in PDB entry: 4ZGM, PDB file 4452.pdb and PDB file A747.pdb, with **A** for GLP-1R ECD and **B** for semaglutide.

| ResName | ResID | Chain ID | ResName | ResID | Chain ID |
|---|---|---|---|---|---|
| THR | 29 | A | SER | 18 | B |
| THR | 29 | A | GLU | 21 | B |
| THR | 29 | A | GLY | 22 | B |
| VAL | 30 | A | GLU | 21 | B |
| VAL | 30 | A | GLY | 22 | B |
| VAL | 30 | A | ALA | 25 | B |
| SER | 31 | A | GLU | 21 | B |
| SER | 31 | A | ALA | 25 | B |
| LEU | 32 | A | PHE | 28 | B |
| LEU | 32 | A | GLU | 21 | B |
| LEU | 32 | A | ALA | 25 | B |
| LEU | 32 | A | ALA | 24 | B |
| THR | 35 | A | PHE | 28 | B |
| THR | 35 | A | ALA | 25 | B |
| THR | 35 | A | ILE | 29 | B |
| VAL | 36 | A | PHE | 28 | B |
| TRP | 39 | A | PHE | 28 | B |
| TRP | 39 | A | LEU | 32 | B |
| TRP | 39 | A | ARG | 36 | B |
| ARG | 43 | A | ARG | 36 | B |
| ASP | 67 | A | GLY | 35 | B |
| ASP | 67 | A | LEU | 32 | B |
| GLU | 68 | A | LEU | 32 | B |
| GLU | 68 | A | GLY | 35 | B |
| GLU | 68 | A | ARG | 36 | B |
| TYR | 69 | A | LEU | 32 | B |
| TYR | 69 | A | ILE | 29 | B |
| TYR | 69 | A | VAL | 33 | B |
| TYR | 88 | A | PHE | 28 | B |
| TYR | 88 | A | LEU | 32 | B |
| TYR | 88 | A | ILE | 29 | B |
| LEU | 89 | A | ILE | 29 | B |
| PRO | 90 | A | ALA | 25 | B |
| PRO | 90 | A | ILE | 29 | B |
| PRO | 90 | A | LYS | 26 | B |
| TRP | 91 | A | LYS | 26 | B |
| TRP | 91 | A | ILE | 29 | B |
| ARG | 121 | A | ARG | 34 | B |
| ARG | 121 | A | LEU | 32 | B |
| ARG | 121 | A | VAL | 33 | B |
| LEU | 123 | A | VAL | 33 | B |
| GLU | 128 | A | LYS | 26 | B |

Table 4: Intermolecular residue contact analysis of PDB entry: 4ZGM [5] by Prodigy [3, 4]. In this table, **ResName** represents the three-letter name of the amino acid residue, **ResID** represents the identification number of the amino acid residue in PDB entry: 4ZGM, **ChainID** specifies the molecular entity in PDB entry: 4ZGM, with **A** for GLP-1R ECD and **B** for semaglutide. In this study, the intermolecular residue contacts at the interface between two molecule entities are defined within the threshold distance of 5.0 Å, as used by the Prodigy [3, 4] server.

| ResName | ResID | Chain ID | ResName | ResID | Chain ID |
|---|---|---|---|---|---|
| THR | 1 | A | GLU | 112 | B |
| THR | 1 | A | SER | 109 | B |
| THR | 1 | A | GLY | 113 | B |
| VAL | 2 | A | GLU | 112 | B |
| VAL | 2 | A | ALA | 116 | B |
| VAL | 2 | A | GLY | 113 | B |
| SER | 3 | A | ALA | 116 | B |
| SER | 3 | A | GLU | 112 | B |
| LEU | 4 | A | ALA | 116 | B |
| LEU | 4 | A | GLU | 112 | B |
| LEU | 4 | A | ALA | 115 | B |
| LEU | 4 | A | PHE | 119 | B |
| THR | 7 | A | ALA | 116 | B |
| THR | 7 | A | PHE | 119 | B |
| VAL | 8 | A | PHE | 119 | B |
| TRP | 11 | A | ARG | 127 | B |
| TRP | 11 | A | PHE | 119 | B |
| TRP | 11 | A | ARG | 123 | B |
| ARG | 15 | A | ARG | 127 | B |
| PHE | 38 | A | ARG | 123 | B |
| ASP | 39 | A | GLY | 126 | B |
| ASP | 39 | A | ARG | 123 | B |
| GLU | 40 | A | ARG | 123 | B |
| GLU | 40 | A | GLY | 126 | B |
| GLU | 40 | A | ARG | 127 | B |
| TYR | 41 | A | GLN | 120 | B |
| TYR | 41 | A | ARG | 123 | B |
| TYR | 41 | A | ARG | 124 | B |
| TYR | 60 | A | GLN | 120 | B |
| TYR | 60 | A | PHE | 119 | B |
| TYR | 60 | A | ARG | 123 | B |
| LEU | 61 | A | GLN | 120 | B |
| PRO | 62 | A | GLN | 120 | B |
| PRO | 62 | A | LYS | 117 | B |
| PRO | 62 | A | ALA | 116 | B |
| TRP | 63 | A | GLN | 120 | B |
| TRP | 63 | A | LYS | 117 | B |
| ARG | 93 | A | ARG | 123 | B |
| ARG | 93 | A | ARG | 124 | B |
| ARG | 93 | A | ALA | 125 | B |
| LEU | 95 | A | GLN | 120 | B |
| LEU | 95 | A | ARG | 124 | B |
| GLU | 99 | A | ARG | 124 | B |
| GLU | 100 | A | LYS | 117 | B |

Table 5: Intermolecular residue contact analysis of PDB file A747.pdb by Prodigy [3, 4]. In this table, **ResName** represents the three-letter name of the amino acid residue, **ResID** represents the identification number of the amino acid residue in PDB file A747.pdb, **ChainID** specifies the molecular entity in PDB file A747.pdb, with **A** for GLP-1R ECD and **B** for semaglutide. In this study, the intermolecular residue contacts at the interface between two molecule entities are defined within the threshold distance of 5.0 Å, as used by the Prodigy [3, 4] server.

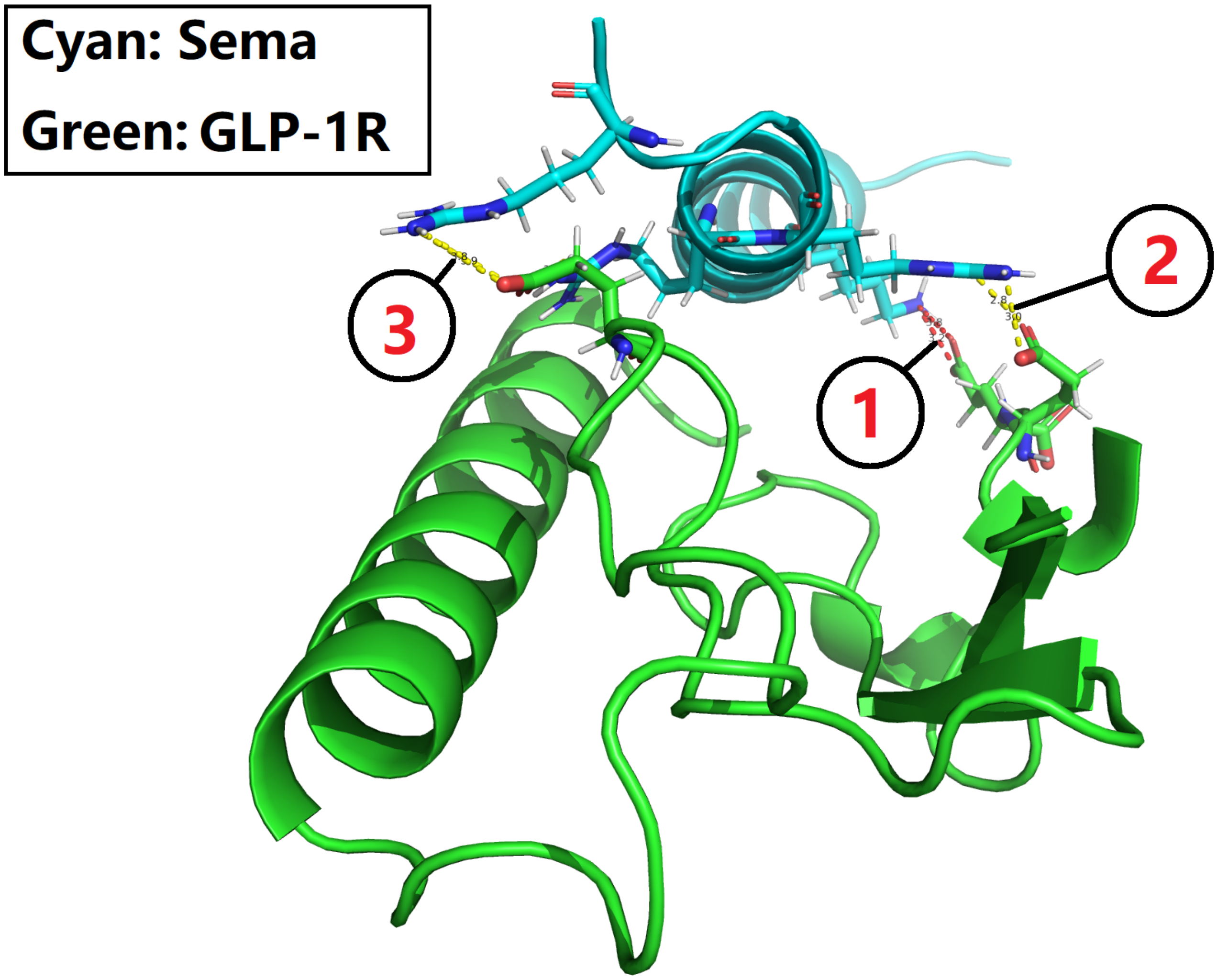


Figure 1: A sideview of the complex structure of GLP-1R ECD and the peptide backbone of semaglutide mutant A747. In this figure, the three red numbers (1, 2 and 3) within black circles represent three sets of interfacial salt bridges (red and yellow dotted lines) of GLP-1R ECD and the peptide backbone of semaglutide mutant A747. These interfacial salt bridges constitute an electrostatic scaffold for the design of next-generation GLP-1R agonists with improved GLP-1R ECD affinity to enhance GLP-1R activation. This figure is prepared with PyMol [6] with supplementary file **A747.pdb**, with number 1 and red dotted lines representing the interfacial salt bridge identified in both PDB entry 4ZGM [5, 7] and **A747.pdb**, and numbers 2 and 3 and yellow dotted lines representing newly established interfacial salt bridges identified in **A747.pdb** but not identified in PDB entry 4ZGM [5, 7].

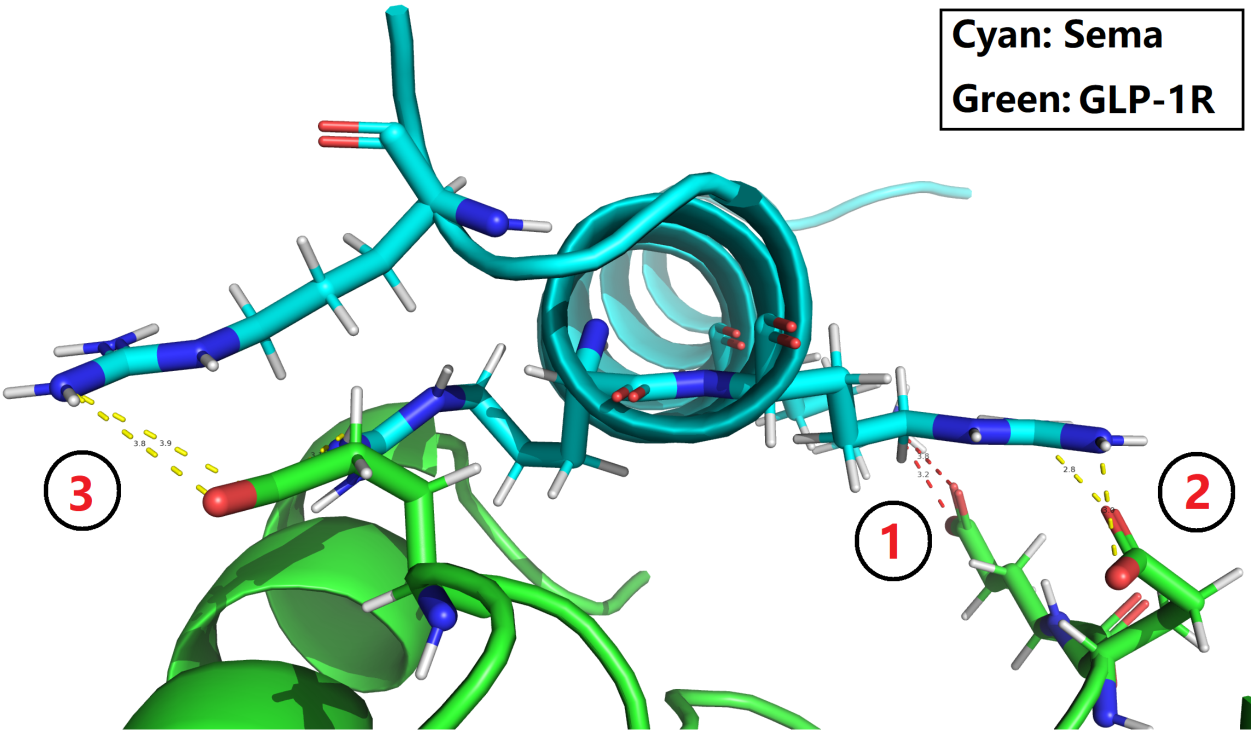


Figure 2: A zoomed-in sideview of the complex structure of GLP-1R ECD and the peptide backbone of semaglutide mutant A747. In this figure, the three red numbers (1, 2 and 3) within black circles represent three sets of interfacial salt bridges (red and yellow dotted lines) of GLP-1R ECD and the peptide backbone of semaglutide mutant A747. These interfacial salt bridges constitute an electrostatic scaffold for the design of next-generation GLP-1R agonists with improved GLP-1R ECD affinity to enhance GLP-1R activation. This figure is prepared with PyMol [6] with supplementary file **A747.pdb**, the details of the three sets of interfacial salt bridges are included in both Table 5 in the main manuscript, and the **Graphical Abstract** of this manuscript, and also in three following figures, i.e., Figures 3, 4 and 5 in this supplementary file.

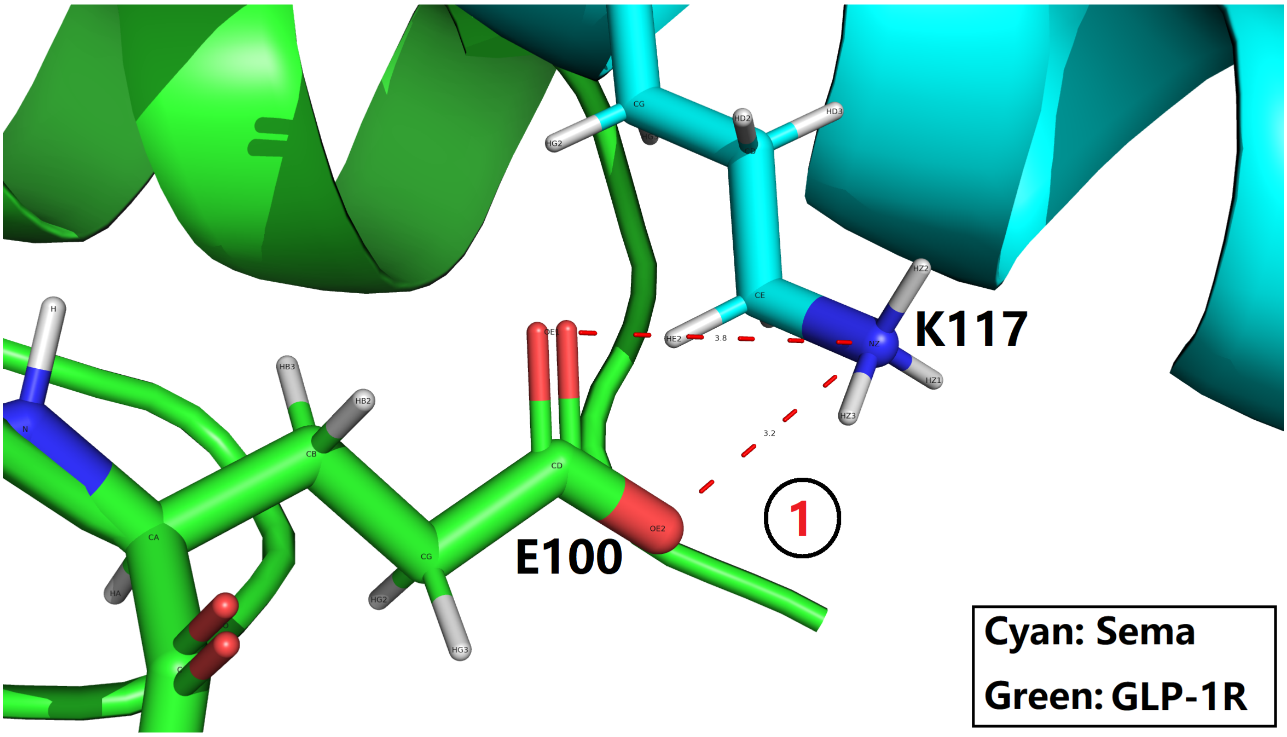


Figure 3: The first (number 1 in Figures 1 and 2) set of interfacial salt bridges (red dotted lines) of GLP-1R ECD and the peptide backbone of semaglutide mutant A747, the details of which are included in Table 5 in the main manuscript, and also included in the **Graphical Abstract** of this manuscript. This figure is prepared with PyMol [6] with supplementary file **A747.pdb**, where K117 and E100 represent Lys117 of semaglutide mutant A747 and Glu100 of GLP-1R ECD, respectively.

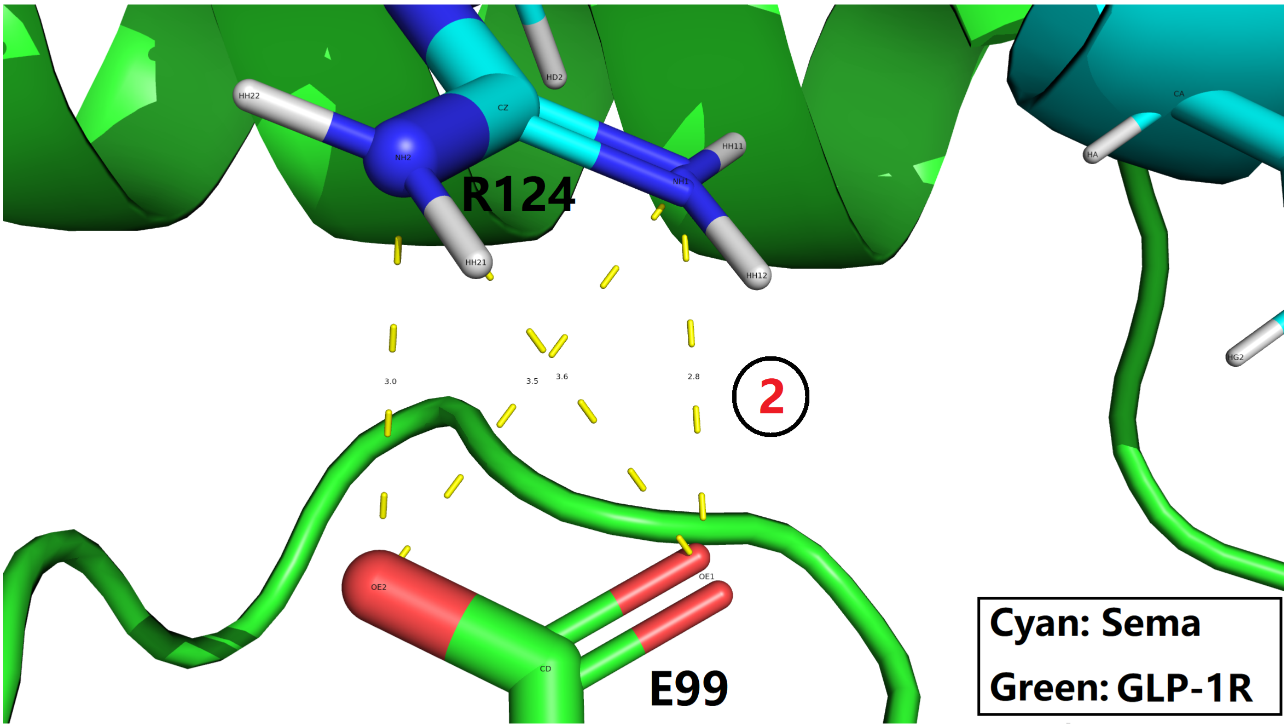


Figure 4: The second (number 2 in Figures 1 and 2) set of interfacial salt bridges (yellow dotted lines) of GLP-1R ECD and the peptide backbone of semaglutide mutant A747, the details of which are included in Table 5 in the main manuscript, and also included in the **Graphical Abstract** of this manuscript. This figure is prepared with PyMol [6] with supplementary file **A747.pdb**, where R124 and E99 represent Arg124 of semaglutide mutant A747 and Glu99 of GLP-1R ECD, respectively.

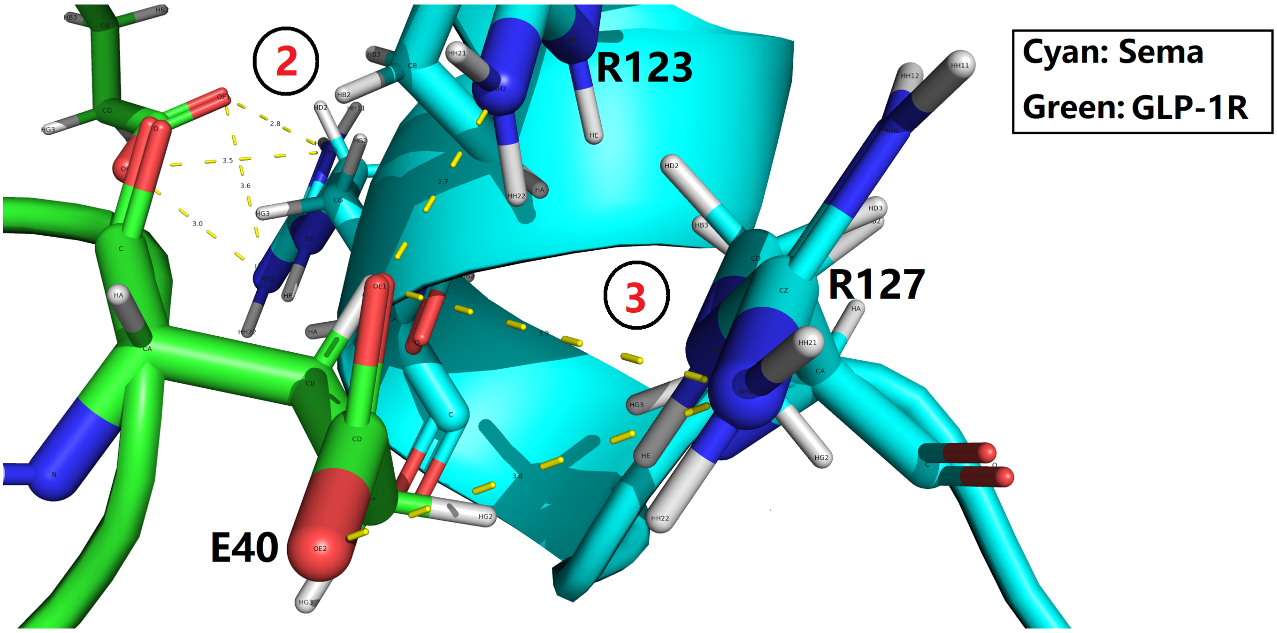


Figure 5: The third (number 3 in Figures 1 and 2) set of interfacial salt bridges (yellow dotted lines) of GLP-1R ECD and the peptide backbone of semaglutide mutant A747, the details of which are included in Table 5 in the main manuscript, and also included in the **Graphical Abstract** of this manuscript. This figure is prepared with PyMol [6] with supplementary file **A747.pdb**, where R123, R127 and E40 represent Arg123 and Arg127 of semaglutide mutant A747, and Glu40 of GLP-1R ECD, respectively.

### References

[1] Li W. In Silico Generation of Structural and Intermolecular Binding Affinity Data with Reasonable Accuracy: Expanding Horizons in Drug Discovery and Design. 2024;doi:10.20944/preprints202405.1739.v1.

[2] Webb B, Sali A. Protein Structure Modeling with MODELLER. In: Methods in Molecular Biology. Springer US; 2020. p. 239–255.

[3] Vangone A, Bonvin AM. Contacts-based prediction of binding affinity in protein-protein complexes. eLife. 2015;4.

[4] Xue LC, Rodrigues JP, Kastritis PL, Bonvin AM, Vangone A. PRODIGY: a web server for predicting the binding affinity of protein-protein complexes. Bioinformatics. 2016; p. btw514.

[5] Lau J, Bloch P, Schäffer L, Pettersson I, Spetzler J, Kofoed J, et al. Discovery of the Once-Weekly Glucagon-Like Peptide-1 (GLP-1) Analogue Semaglutide. Journal of Medicinal Chemistry. 2015;58(18):7370–7380.

[6] DeLano WL. Pymol: An open-source molecular graphics tool. CCP4 Newsletter On Protein Crystallography. 2002;40:82–92.

[7] Reedtz-Runge S. Crystal structure of Semaglutide peptide backbone in complex with the GLP-1 receptor extracellular domain; 2015.
